# Blunted Diurnal Firing in Lateral Habenula Projections to Dorsal Raphe Nucleus and Delayed Photoentrainment in Stress-Susceptible Mice

**DOI:** 10.1101/2020.03.19.998732

**Authors:** He Liu, Ashutosh Rastogi, Merima Sabanovic, Aisha Darwish Alhammadi, Qing Xu, Lihua Guo, Jun-Li Cao, Hongxing Zhang, Priyam Narain, Hala Aqel, Vongai Mlambo, Rachid Rezgui, Basma Radwan, Dipesh Chaudhury

**Affiliations:** The Division of Science, The New York University Abu Dhabi, Abu Dhabi, United Arab Emirates; Department of Anesthesiology, The Affiliated Hospital of Xuzhou Medical University, Xuzhou, China; Jiangsu Province Key Laboratory of Anesthesiology & Jiangsu Province Key Laboratory of Anesthesia and Analgesia Application Technology, The Xuzhou Medical University, Xuzhou, China

**Keywords:** Stress, Lateral Habenula, Dorsal Raphe Nucleus, Diurnal, Firing-Rate, Photoentrainment

## Abstract

Daily rhythms are disrupted in patients suffering from mood disorders. The lateral habenula (LHb) and dorsal raphe nucleus (DRN) contribute to circadian timekeeping and regulate mood. Thus, pathophysiology in these nuclei may be responsible for aberrations in daily rhythms during mood disorders. Using the chronic social defeat stress (CSDS) paradigm and *in-vitro* slice electrophysiology we measured the effects of stress on diurnal rhythms in firing of LHb cells projecting to the DRN (cells^LHb→DRN^) and DRN cells alone. We also performed optogenetic experiments to investigate if increased firing in cells^LHb→DRN^ during exposure to subthreshold social defeat stress (SSDS), induces stress-susceptibility. Last we investigated whether exposure to CSDS affected the ability of mice to phototentrain to a new LD cycle. The cells^LHb→DRN^ and DRN cells alone of stress-susceptible mice express greater blunted diurnal firing compared to stress-naive (control) and stress-resilient mice. Day-time optogenetic activation of cells^LHb→DRN^ during SSDS induces stress-susceptibility which shows the direct correlation between increased activity in this circuit and putative mood disorders. Finally, we found that stress-susceptible mice are slower, while stress-resilient mice are faster, at photoentraining to a new LD cycle. Our findings suggest that CSDS induces blunted daily rhythms in firing in cells^LHb→DRN^ and slow rate of photoentrainment in susceptible-mice. In contrast, resilient-mice may undergo homeostatic adaptations that maintain daily rhythms in firing in cells^LHb→DRN^ and also show rapid photoentrainment to a new LD-cycle.

## Introduction

Mood disorders are associated with abnormalities in circadian rhythms in physiology and behaviour. The LHb and DRN regulate diverse behaviours associated with mood, including cognition, reward and sleep-wake cycle(1, 2). Moreover, observations that the LHb and DRN are connected to the suprachiasmatic nucleus (SCN), the master circadian clock, and that both nuclei exhibit diurnal rhythms in clock genes expression and neuronal activity(1, 3) highlight the possible evolutionarily significant relationship between circuits that regulate motivational behaviours and the daily rhythms of such behaviours(2). It is hypothesized that pathophysiological changes in these circuits may be responsible for the pathogenesis of psychiatric disorders and associated changes in behavioral rhythms(2, 4).

Given the overlapping influence of the LHb and DRN on circadian rhythmicity and mood disorders, we assessed whether exposure to chronic social defeat stress (CSDS) affects: (i) diurnal rhythmic activity in LHb cells projecting to the DRN (Cells^LHb→DRN^) and (ii) the ability of mice to re-entrain to a new photoperiod. We demonstrate that cells^LHb→DRN^ of stress-susceptible mice exhibit blunted daily rhythms in neural activity. Specifically, the firing rate was elevated in both, the day and night, in susceptible-mice while resilient and stress-naïve mice display robust diurnal rhythms in firing with high activity in night and low in daytime. The blunted rhythms in firing-rate in stress-susceptible mice is similar to observations that patients with mood disorders exhibit blunted daily rhythms in various physiological measures(5, 6). In addition to pathophysiological changes in firing in the cells^LHb→DRN^, susceptible-mice also exhibit delayed rate of re-entrainment to a new light cycle while resilient mice were faster at adapting to the new light cycle. Our findings suggest that chronic social stress induces pathophysiological firing in cells^LHb→DRN^ of susceptible-mice that leads to blunted diurnal activity in this circuit. Moreover, the behavioral locomotor data suggests that molecular and cellular processes responsible for acute photoentrainment seems slower at responding to a new light stimulus in susceptible-mice while homeostatic adaptations in resilient mice increase their efficiency at adapting to the new light cycle.

## Results

### Increased Day-Time Firing in LHb Cells of Susceptible-Mice

Though chronic stress leads to increased activity in the LHb(7), it is unclear whether the LHb of mice resilient or susceptible to CSDS express the same increase in firing. To explore this, we performed *in-vivo*, extracellular single-unit recordings in daytime in anesthetized mice that had been undergone CSDS (**Figure 1A-B** and **Supplemental Figure 1A-B**). The spontaneous firing-rate was significantly higher in susceptible-mice than control (stress-naïve) and stress-resilient-mice (**Figure. 1C**), which suggests that increased day-time activity in the LHb cells encode for susceptibility to CSDS. Though not significant there is a slight trend showing a negative correlation between firing rate and social interaction ratio (**Supplemental Figure 1C**).

**Figure 1.**
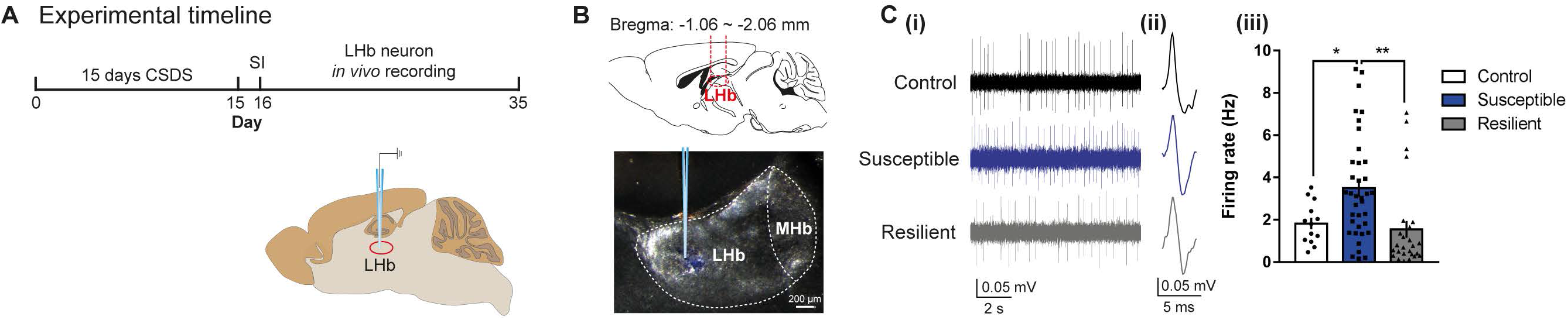
Increased Day-Time Firing in LHb Cells of Susceptible-Mice. **(A)** Experimental timeline of Chronic Social Defeat Stress (CSDS) paradigm, social interaction (SI), and *in-vivo* single-unit recording in the day-time. **(B)** Schematic showing anatomical site of LHb (top); and example site of glass electrode recording in the LHb (bottom). **(C) (i)** Example traces, **(ii)** typical action potentials of recorded LHb cells of control (top), susceptible (middle) and resilient mice (bottom) after SI test. **(iii)** Increased firing-rate in susceptible-mice (*F*_2,73_ = 7.195, *P* = 0.0014; n = 4 mice/group, n = 13-37 cells/group). Abbreviations: LHb, lateral habenula: MHb, medial habenula. Error bars: mean ± SEM.

### Blunted Rhythms in Spontaneous Firing in Cells^LHb→DRN^ of Susceptible-Mice

The majority of LHb cells express peak firing, and increased c-Fos expression during the night(8, 9). Moreover, the LHb has reciprocal connections with the DRN(7), a region that modulates sleep-wake states(10) and mood(11). Thus, we wondered whether exposure to CSDS affected diurnal rhythmic activity in cells^LHb→DRN^. Immunohistochemical labelling showed that approximately 5%<u>±</u>0.1% of cells from the medial and lateral portion of the LHb (LHbM and LHbL) project to the DRN (**Figure 2-C**). Next, we investigated the diurnal firing properties of cells^LHb→DRN^ in mice exposed to CSDS (**Figures 2D-E**). The SI scores for control, resilient and susceptible-mice were comparable for animals where electrophysiological recordings were performed in the day or the night (**Supplemental Figures 2A-B**). After the SI test, mice were euthanized in the day (ZT1) or the night (ZT13) after which slice electrophysical recordings were conducted in fluorescently labelled cells^LHb→DRN^ (**Supplemental Figure 2C**). Day-time spontaneous firing-rate of cells^LHb→DRN^ of susceptible-mice was significantly higher (**Figure 2D**). In contrast, there was no difference in night-time spontaneous firing-rate in cells^LHb→DRN^ between control, resilient and susceptible-mice (**Figure 2E**). Moreover, we show a significant negative correlation between firing rate and social interaction ratio during the day but not at night (**Figure 2F-G**). Re-plotting the data of the spontaneous activity in the 1^st^ and 2^nd^ week of recordings showed that control and resilient-mice exhibited robust differences in the pattern of diurnal activity where activity was lower in the day and higher at night, while in susceptible-mice the firing-rate remained elevated in both the day and night leading to a blunted rhythm (**Supplemental Figure 2D**). Since LHb cells exhibit silent, tonic or burst-firing characteristics(12), we investigated whether exposure to stress affected the diurnal firing characteristics in cells^LHb→DRN^. All three behavioral phenotypes exhibited a greater percentage of burst-firing cells at night-time (**Figure 2H**). Intrinsic membrane properties also show diurnal rhythmicity such that during the day, cells^LHb→DRN^ of susceptible-mice were significantly more excitable as evident by increased firing in response to current injection (**Supplemental Figure 3A-top**). In accord with increased evoked-excitability, susceptible-mice displayed reduced threshold to induce the first spike (rheobase) in response to current injection and increased membrane input-resistance in the steady-state current-voltage (I-V) relationship (**Supplemental Figure 3B-C-top**). In contrast, at night there was no difference in evoked excitability, rheobase or input resistance between any of the three behavioral groups (**Supplemental Figures 3A-C-bottom**).

**Figure 2.**
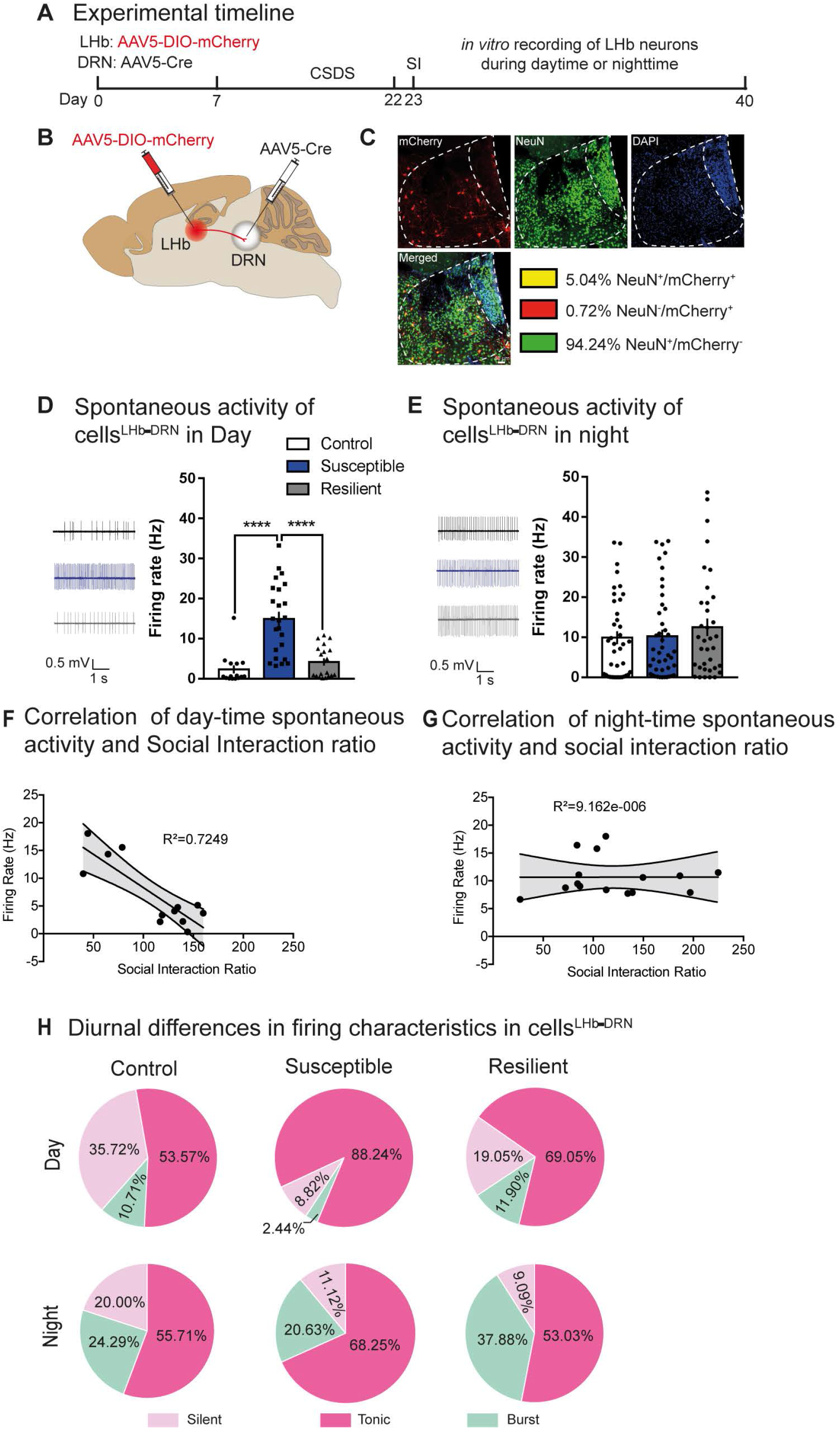
Blunted Rhythms in Spontaneous Firing in Cells^LHb→DRN^ of Susceptible-Mice. **(A)** Experimental timeline of viral surgeries to label cells^LHb-DRN^, chronic social defeat stress (CSDS), social interaction (SI) test and *in-vitro* slice electrophysiology. **(B)** Schematic showing surgeries for viral injections to specifically label cells^LHb-DRN^. **(C)** Confocal images showing co-expression of mCherry, NeuN and DAPI within LH; and quantified data showing mCherry-expressing neurons (5% ± 0.1% LHb neurons project to DRN). (**D)** Sample traces of day-time *in vitro* spontaneous activity in cells^LHb-DRN^ of mice exposed to CSDS (left). Increased day-time spontaneous activity in cells^LHb-DRN^ in susceptible-mice (right) (*F*_2,55_ = 22.68, *P* < 0.0001; n = 15-24 cells from 7 to 11 mice/group). **(E)** Sample traces of night-time *in vitro* spontaneous activity in cells^LHb-DRN^ of mice exposed to CSDS (left). No difference in night-time spontaneous firing in cells^LHb-DRN^ in control, susceptible and resilient-mice (right). (**F-G**) Significant negative correlation between spontaneous activity in cells^LHb-DRN^ and social interaction ratio during the day (R^2^=07249; p<0.05) but not the night. **(H)** Pie charts illustrating the percentage of cells expressing: silent, tonic- or burst-firing in the day-time (top) and night-time (bottom) in labelled cells^LHb-DRN^. Susceptible-mice exhibit notable increased day-time tonic-firing and decrease in silent patterns (*χ2* = 6.947, *P* < 0.05, Chi-square test; n = 28-42 cells from 7 to 11 mice/group). At night, there is no difference in firing patterns between control-, susceptible- and resilient-mice. There is significant increase in burst firing at night compared to day (P < 0.05). Error bars: mean ± SEM.

### CSDS Induces Diurnal Rhythms in I_h_ and K_v_^+^ - Currrents in Cells^LHb→DRN^

Hyperpolarization-activated cation channel (HCN) mediated I_*h*_-current, which have an excitatory drive, undergo diurnal rhythmic expression and are differentially regulated in susceptible and resilient-mice(13, 14). We tested whether the diurnal differences in excitability in cells^LHb→DRN^ of susceptible and resilient-mice was due to changes in the strength of the I_*h*_-currents. Our observations that I_*h*_-currents of control-mice are equally lower in the day and the night (**Figure 3A-B**) suggests that these channels do not drive the diurnal difference in firing in this circuit in non-stressed conditions. However, exposure to chronic stress triggers rhythmic expression of HCN-channels since both susceptible and resilient-mice show larger I_*h*_-currents in the day than night (**Figure 3A-B**). Moreover, it reflects the diurnal differences in evoked-excitability (**Supplemental Figure 3A**).

**Figure 3.**
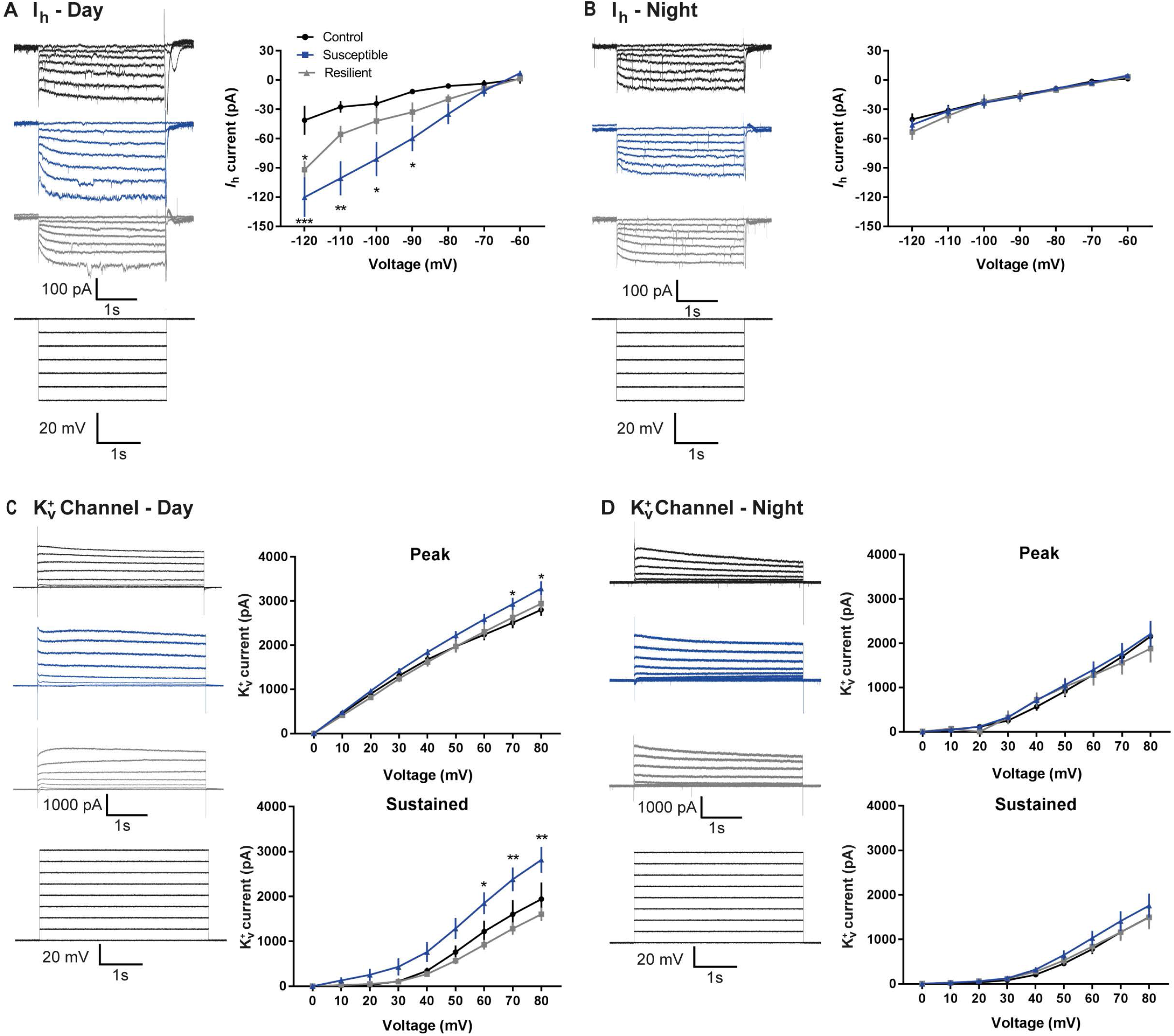
Chronic Social Stress Induces Diurnal Rhythms in I_h_ and K_v_^+^ -Currrents in Cells^LHb→DRN^. **(A)** Day-time representative traces (left) and statistical data of I_h_-currents in response to incremental steps in voltage injections (right). Resilient- and susceptible-mice display significant increase in I_h_- current compared to control (*F*_2,252_ = 23.29, *P* < 0.0001; n = 10 to 14 cells from 7 to11 mice per group). **(B)** Night-time representative traces (left) and statistical data of I_h_-currents in response to incremental steps in voltage injections (right). There was no difference in current between any of the phenotypes. **(C)** Day-time representative traces (left) and statistical data of isolated K_v_^+^ channel–mediated currents (right). Susceptible-mice display significant increase in peak (*F*_2,216_ = 9.94, *P* < 0.0001; n = 6 to 11 cells from 7 to 8 mice per group; top) and sustained (*F*_2,216_ = 24.97, *P* < 0.0001; n = 6 to 11 cells from 7 to 8 mice per group; bottom) phases of K_v_^+^-currents. (**D)** Night-time representative traces (left) and statistical data of isolated K_v_^+^ channel–mediated currents (right). There is no difference in peak and sustained phases of K_v_^+^-currents among the three phenotypes. Error bars: mean ± SEM.

Voltage-gated potassium (K_v_^+^)-channels, which regulate cell-excitability, are affected by stress-exposure and undergo diurnal rhythmic expression(14–16). Thus, we investigated whether these currents are responsible for the daily differences in firing in cells^LHb→DRN^. Since subtypes of K_v_^+^-channels exhibit different kinetics, we analysed both peak (fast-kinetics) and sustained (slow-kinetics) K_v_^+^-currents in the day and night following CSDS. We found that peak K_v_^+^-currents are larger in the day in all behavioral groups with susceptible-mice showing the largest day-time peak current than night (**Figure 3C-D-top**). This shows that fast-kinetics K_v_^+^-channels undergo daily rhythmic expression in this circuit, irrespective of stress-exposure. In contrast, sustained K_v_^+^-currents do not exhibit daily rhythmicity in control-mice implying that slow-kinetics K_v_^+^-channels may not regulate diurnal changes in firing-rate in cells^LHb→DRN^ in non-stressed conditions (**Figure 3C-D-Bottom**). However, exposure to stress likely induces molecular changes that activate rhythmic expression of slow-kinetic K_v_^+^-channels in susceptible, but not, resilient-mice in daytime (**Figure 3C-bottom**). It is possible that homeostatic adaptations in resilient-mice prevent the stress-induced diurnal expression of these channels. Since the majority of K_v_^+^-currents have an inhibitory drive, our observation that control- and resilient-mice express lower day-time spontaneous-firing may in part be due to the day-time increase in peak K_v_^+^-currents. However, the large day-time peak and sustained K_v_^+^-currents in susceptible-mice was unexpected.

### Day-Time Optical Activation of Cells^LHb→DRN^ During Weak Social Stress Induces Susceptibility

Our observation that the spontaneous firing-rate in cells^LHb→DRN^ of susceptible-mice is elevated in the day and the night lead us to speculate that blunted diurnal rhythmic firing in this circuit may represent pathophysiology leading to mood disorders. We performed *in-vivo* optogenetics experiments to directly correlate the link between increased day-time firing in cells^LHb→DRN^ and stress-susceptibility. We first validated that optical stimulation with blue light (473nm) induces action potentials and inward-currents (**Supplemental Figure 4A-B-left**). Frequency-response curves of membrane excitability showed fidelity up to 20Hz following optical stimulation (**Supplemental Figure 4B-right**). Next, we performed in-vivo optogenetic experiments where mice were exposed to a SSDS paradigm during which the ChR2 expressing cells^LHb→DRN^ were optically stimulated for 20-mins after social stress (**Figure 4A**, **Supplemental Figure 4C-D**). Mice that undergo SSDS do not typically exhibit social avoidance or other stress-susceptible behaviours but are more vulnerable to subsequent stress(17, 18). Optical stimulation of cells^LHb→DRN^ during SSDS (SSD-ChR2) induced stress-susceptibility as shown by the significant decrease in SI-ratio compared to mice exposed to SSDS alone (SSD-mCherry) or non-SSDS-exposed mice that underwent optical stimulation of cells^LHb→DRN^ (Non-SSD-ChR2: **Figure 4B**, **Supplemental Figure 4E**). Our findings that pro-susceptibility was only observed when cells^LHb→DRN^ were optically stimulated during SSDS suggests that the pro-stress effects of increased firing in this circuit is context specific. Moreover, the effect is long-lasting because SSD-ChR2 mice, but not non-stressed SSD-mCherry and Non-SSD-Chr2 groups, continue to show stress-susceptibility because they express anhedonic traits (decreased sucrose preference) 36-days after the end of the SSDS and optogenetic stimulation (**Figure 4C**).

**Figure 4.**
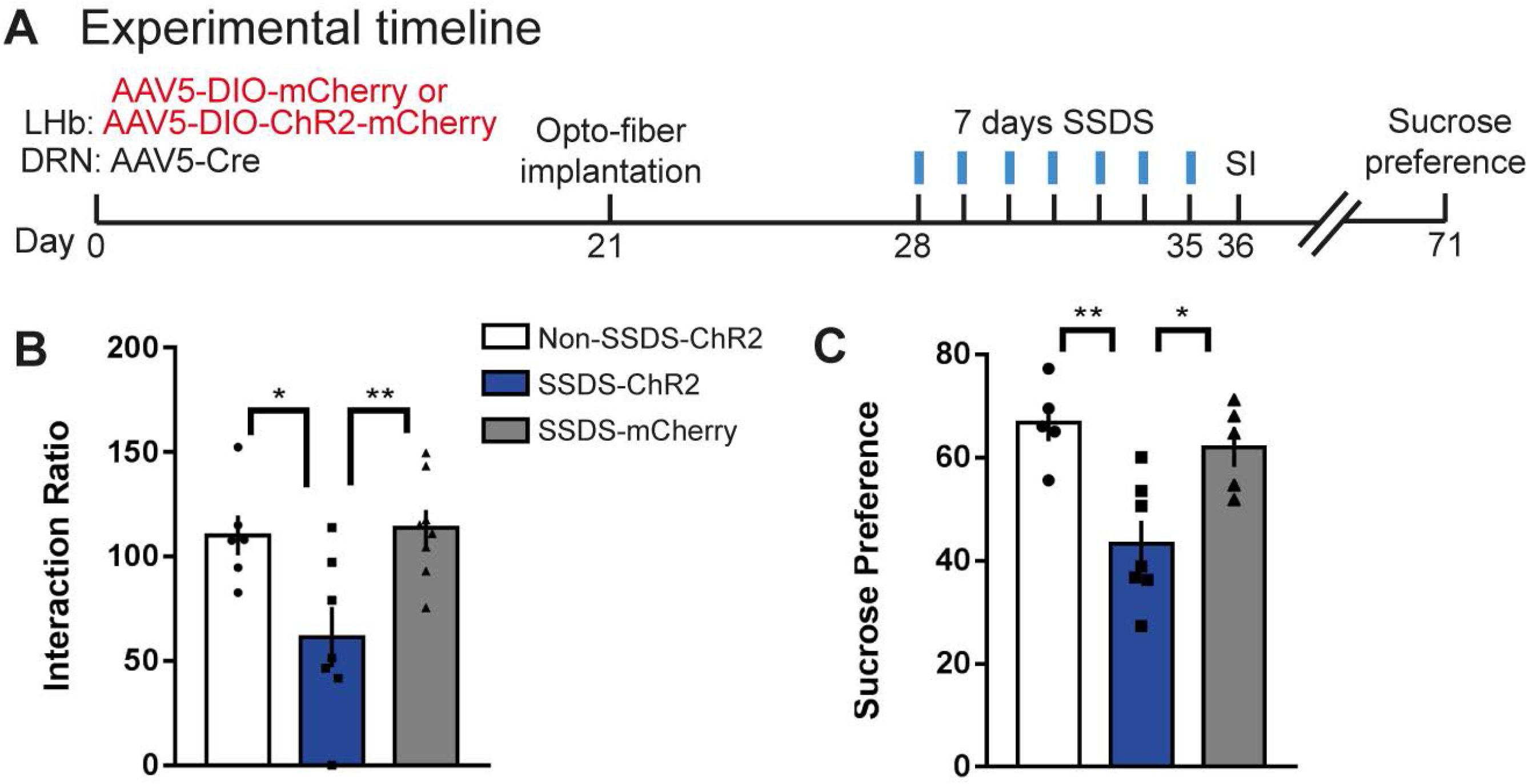
Day-Time Optical Activation of Cells^LHb→DRN^ During Weak Social Stress Induces Susceptibility. **(A)** Experimental timeline of viral surgeries, opto-fiber implantation, repeated high frequency optical stimulation patterns and behavioral tests. (**B)** In the presence of a CD1 social target (non-aggressor), SSD-ChR2 mice display increased stress-susceptibility as measured by decreased SI ratio compared with Non-SSD-ChR2 and SSD-mCherry mice, (*F*_2,18_ = 6.905, *P* < 0.01; n = 6-8 mice/group). (**C)** The SSD-ChR2 mice display anhedonia as measured by reduction in 1% sucrose intake over 3 hrs in the sucrose preference test (*F*_2,14_ =9.805, *P* < 0.01; n = 5-7 mice/group; right). Error bars: mean ± SEM.

### Mice Susceptible or Resilient to Chronic Social Stress Exhibit Different Rates of Photoentrainment

To assess the effects of a chronic social stressor on diurnal and circadian rhythms, we measured locomotor wheel-running activity in stress-naïve, susceptible and resilient-mice for 7-days during a standard 12hr:12hr LD-cycle followed by 7-days of DD (**Figure 5A**, **Supplemental Figure 5A**). Exposure to CSDS did not affect endogenous rhythms since there was no difference in the phase of activity onset in DD relative to LD between any of the groups (**Supplemental Figures 5B-D**). Moreover, there was no difference in the free-running period in DD and amount of activity in LD and DD (**Supplemental Figures 5E-F**). We also measured photoentrainment in stress-exposed mice challenged by a 6-hr advance in LD cycle. Further detailed analysis of behavioural rhythms showed that Intradaily variability was lower in stress exposed mice compared to stress-naïve groups while there was no difference for interdaily stability between the three phenotypes (Supplemental Figures 5G-J). Behavioral wheel running activity was measured for 7-days in standard LD cycle followed by activity measurement for 7-days after a single 6-hr advance in light onset (**Figures 5A-B**). Susceptible-mice exhibited significantly slower initial rate of photoentrainment while resilient-mice were significantly quicker at fully re-entraining to the new photoperiod (**Figures 5C-D**).

**Figure 5.**
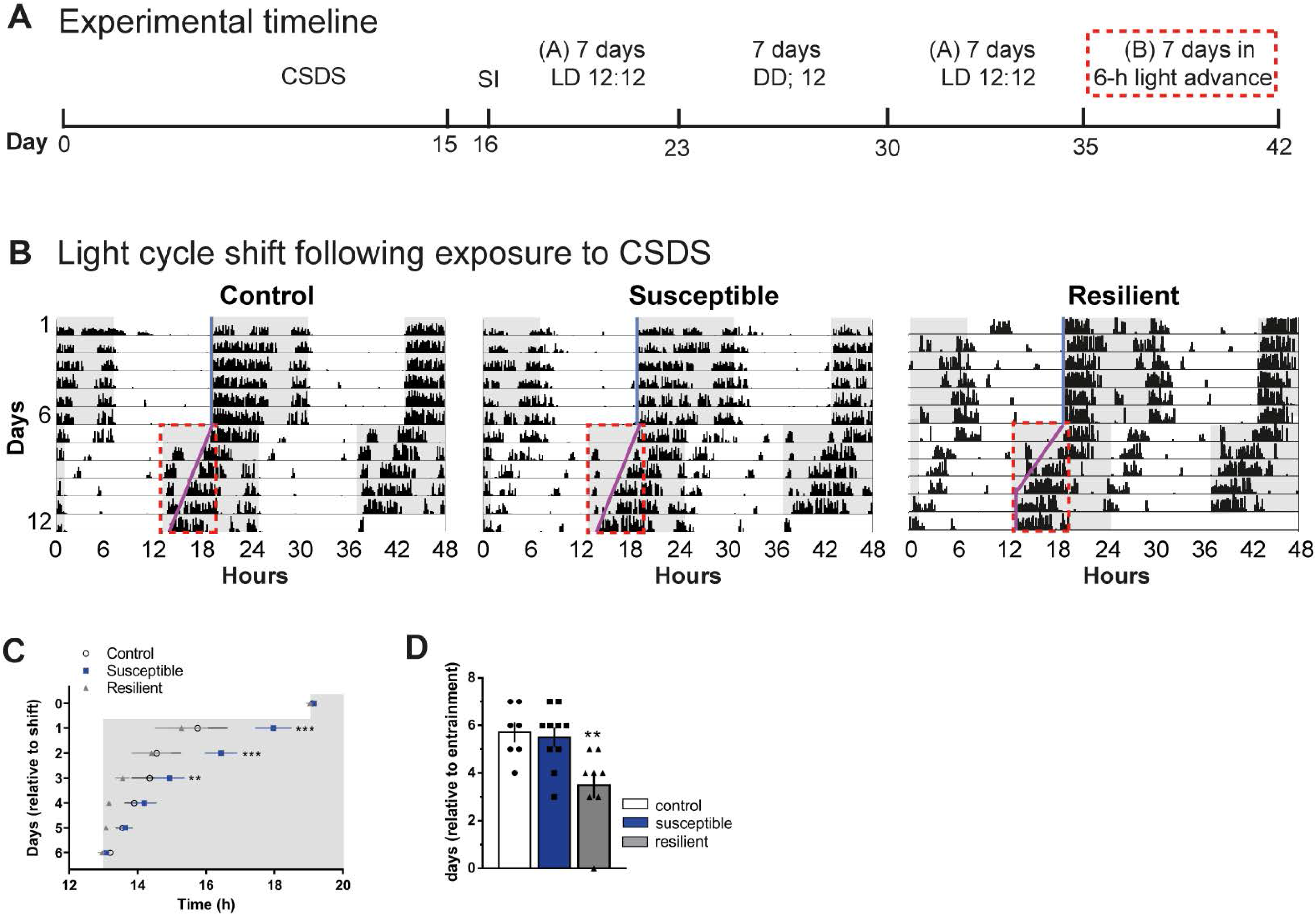
Mice Susceptible or Resilient to Chronic Social Stress Exhibit Different Rates of Photoentrainment. **(A)** Experimental timeline (A - normal light cycle: Lights ON: 07:00; Lights OFF: 19:00; B - new light cycle following 6-hrs advance: Lights ON: 13:00; Lights OFF: 01:00), red box denotes 6-hrs lights advance in new LD cycle. **(B)** Representative double-plotted actograms of activity before and after 6-hr shift in LD cycle. Gray background indicating dark phase of the LD cycle. Light blue and pink lines representing extended regression line derived by onset of activity under normal and new LD cycles, respectively. Dotted red lines outlining the data of activity onsets shown in (c). **(C)** Susceptible-mice display significantly slower initial-rate of photoentrainment in the new LD cycle (*F*_2,25_= 10.81, *P* < 0.001; n = 7-10 mice/group). **(D)** Resilient-mice took significantly less number of days to re-entrained in the new LD cycle (*F*_2,22_ = 6.587, *P* < 0.05). Error bars: mean ± SEM.

### Blunted Rhythms in Spontaneous Firing in DRN Cells of Stress-Susceptible Mice

We next investigated spontaneous firing in DRN in the day and the night (**Figure 6A**). During the day there is a bimodal response in CSDS exposed mice, where spontaneous firing in DRN cells is lower in susceptible-, and higher in resilient-mice compared to control groups, while at night there was no difference (**Figures 6B-C**). Furthermore, we show a significant positive correlation between firing rate and social interaction ratio during the day but not at night (**Figure 6D-E**). Comparison of diurnal rhythms in firing shows that control and resilient-mice tend to express higher relative activity in the day while susceptible-mice show blunted rhythms where activity is equally low in the day and the night (**Supplemental Figure 6**).

**Figure 6.**
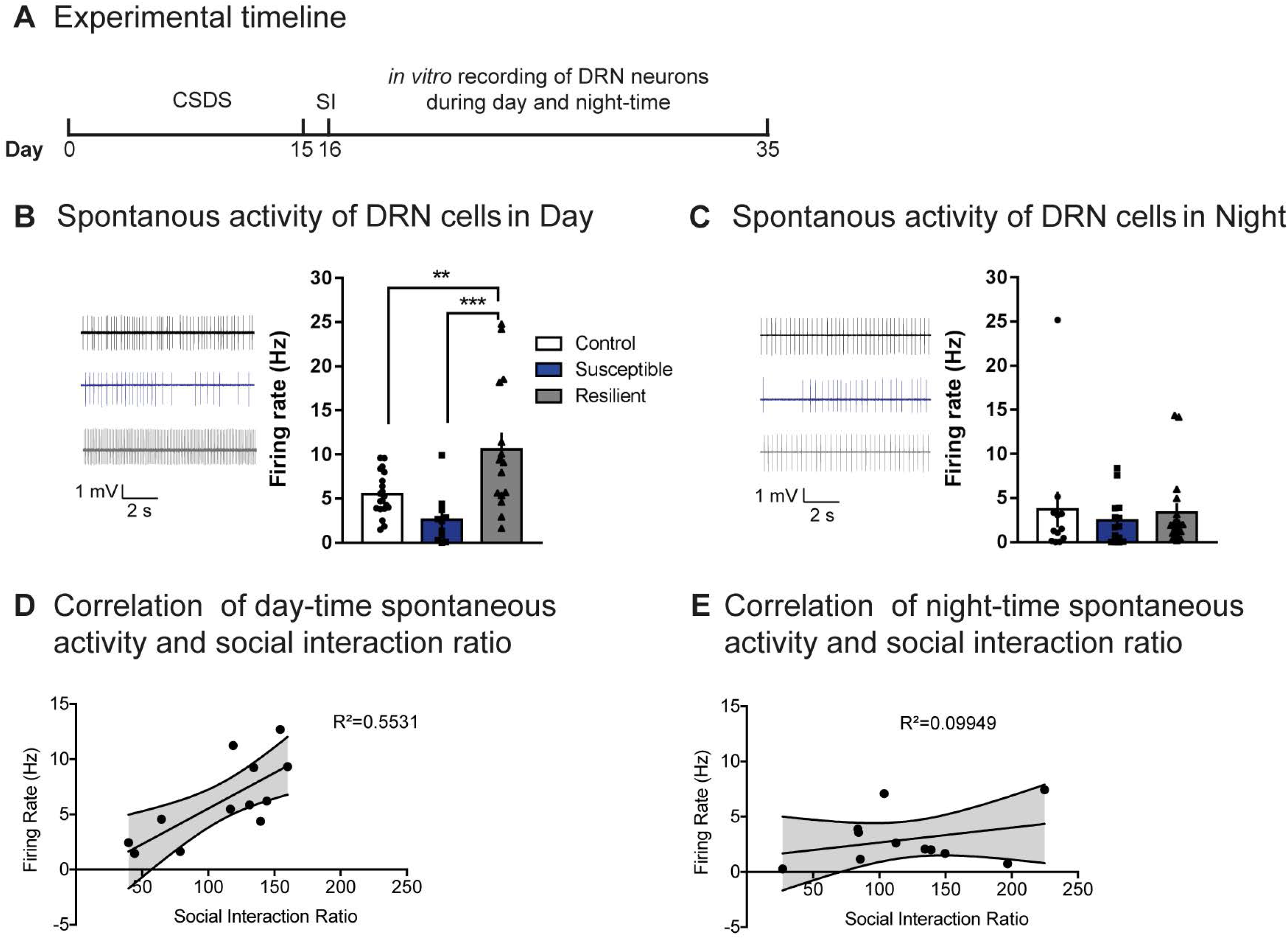
Blunted Rhythms in Spontaneous Firing in DRN Cells of Stress-Susceptible Mice. **(A)** Experimental timeline of CSDS paradigm, SI test, and *in-vitro* recordings. **(B-C)** Sample traces of spontaneous firing of DRN cells in day (B) and in night (C) in control- (top), susceptible- (middle) and resilient-mice (bottom). (B) Susceptible-mice exhibit decreased day-time spontaneous firing in DRN cells while resilient-mice exhibit increased day-time spontaneous firing (*F*_2,43_ = 9.793, *P* < 0.001; n = 11-20 cells from 7 to 11 mice/group), (C) No difference in night-time spontaneous firing in DRN cells. (**D-E**) Significant positive correlation between spontaneous activity in DRN cells and social interaction ratio during the day (R^2^=0.5531; p<0.05) but not the night. Error bars: mean ± SEM.

## Discussion

We found that cells^LHb→DRN^ of susceptible-, but not resilient-mice, exhibit pathophysiological firing such that activity is elevated in both the day and the night. The LHb and DRN have direct and indirect connections with the SCN which likely leads to their functional role in integrating mood related behaviour with circadian timing (4, 19),(1). Thus, we predict that that stress induced pathophysiological firing in the loop consisting of the LHb, DRN and SCN (LHb→DRN→SCN→LHb) may be responsible for changes in daily rhythms associated with mood disorders. Stress and the circadian system exhibit 2-way interaction where for example clock proteins CRY 1/2 increase the responsiveness of the HPA-axis(20) while stress-induced increase in glucocorticoids leads to altered clock gene expression in the SCN and regions associated with regulation of mood(21),(22). Stress induces increased firing in the LHb has been shown to activate HPA-axis causing elevated corticosterone release(23). Thus, it is possible that our observations of elevated firing and blunted daily rhythms in cells^LHb→DRN^ of susceptible-mice may be responsible for blunted rhythms in corticosterone observed in rats exposed to chronic mild stress(24). Blunted diurnal rhythmic expression of clock proteins in the SCN in stress-susceptible rodents(25, 26) likely causes blunted rhythmic firing of SCN cells projecting to downstream targets such as the LHb. We speculate that this process may be responsible for our observations of elevated day-night spontaneous firing in cells^LHb→DRN^ of susceptible-mice, leading to blunted diurnal rhythmicity in neural activity while homeostatic adaptations in the SCN and LHb cells of resilient-mice may prevent stress induced blunting of rhythmic activity.

Synaptic inputs and diurnal changes in ion channel expression regulate duration of action potential, control membrane potential and neural excitability(27). We speculated that stress induced disruptions on the daily oscillations of hyperpolarization-activated cation channel (HCN) mediated I_*h*_-currents and voltage-gated potassium (K_v_^+^)-currents in cells^LHb→DRN^ may be responsible for the elevated diurnal, spontaneous firing in susceptible-mice. Specifically, we predicted that low I_*h*_ and high K_v_^+^-currents during the day would drive low firing in control and resilient-mice that would switch at night leading to high firing while elevated I_*h*_ and decreased K^+^-currents at both the day and the night would drive increased diurnal firing in susceptible-mice. Our findings that both I_*h*_- and sustained K_v_^+^-currents are equally low in the day and night of control-mice suggests that these currents likely do not have a functional role in regulating daily rhythms in spontaneous firing in cells^LHb→DRN^ in non-stressed conditions. However, susceptible-mice show rhythmic expression of these currents (high at day, low at night) suggesting that stress induces changes in molecular pathways triggering increased expression of I_*h*_ and slow kinetic K_v_^+^-channels in the day. The discrepancy between robust daily-rhythms in I_*h*_- and K_v_^+^-currents, but not in spontaneous firing, in susceptible-mice may arise because spontaneous firing is a net measure of synaptic inputs and the effect of numerous ion channels that may dilute the individual effects of I_*h*_- and K_v_^+^- currents. Our observation that more cells exhibit bursting activity at night in all behavioral phenotypes may reflect the increased expression of T-type calcium (Ca^2+^)-channels and calcium-activated-K^+^-channels which have been previously shown to regulate bursting activity in the LHb(28). Moreover, increased night-time expression of these channels may be responsible for the decreased absolute evoked excitability since depolarizing currents may induce increased calcium influx, via T-type Ca^2+^-channels, leading to greater activation of Ca^2+^-activated K^+^-channels and periodic hyperpolarization of cell membrane with consequent decreased propensity to fire. Since K_v_^+^-currents typically have an inhibitory drive that decreases the firing-rate of a cell, the day-time increased peak voltage activated (K_v_^+^)-currents in cells^LHb→DRN^ in control and resilient-mice may be partly responsible for the lower activity in these mice. However, the large peak and sustained K_v_^+^-currents in cells^LHb→DRN^ in susceptible-mice is unexpected. Two possible explanations to account for increased firing with increased K_v_^+^-currents are: (i) as shown in the VTA, increased I_*h*_-currents induces compensatory K_v_^+^-currents(14), however these compensatory currents in cells^LHb→DRN^ may be ineffective at lowering the firing-rate possibly due to a larger net increase in the excitatory driving force and (ii) aberrations in clock genes expression upregulates day-time expression of fast activation and inactivation A-type potassium (IKA) channels and delayed rectifier K_v_^+^-channels (DR). Both these channels increase firing-rate by shortening action potential duration. Moreover, these channels undergo daily oscillations within the SCN such that their peak expression is highest in the day when these cells have the highest firing-rate(15, 16). Thus, one intriguing possibility is that the diurnal rhythms in peak K_v_^+^-currents may represent rhythmic expression of the fast dynamics IKA-channels whose day-time expression is further increased in susceptible mice. Furthermore, large day-time sustained K_v_^+^-currents in susceptible-mice may represent a larger proportion of DR channels whose expression peaks in the day that return to basal levels at night.

Our observation that the rates of re-entrainment to a new LD-cycle differ between susceptible and resilient-mice suggest that stress induces different changes in molecular and cellular processes that regulate photoentrainment. For example, susceptible-mice express decreased amounts of the phosphorylated form of glycogen synthase kinase (p-GSK-3β(29) and since photoentrainment requires conversion of p-GSK-3β to GSK3β)(30) aberrations in this cascade may be responsible for the slow initial rate of photoentrainment observed in the susceptible phenotype. Moreover, in the SCN, salt inducible kinase 1 (SIK1) and CREB-regulated transcription coactivator1 (CRTC1) have a functional role in photoentrainment(31). Specifically, night-time light pulses activate CRTC1 and CREB leading to *Per1* and *Sik1* expression. Increased PER starts photoentrainment however SIK1 forms part of a negative feed-back that prevents further *Per1* expression preventing immediate photoentrainment to a new LD cycle. Since stress increases hypothalamic SIK1 expression(32) it is possible that elevated light-induced SIK1 expression in susceptible-mice may be responsible for the slower initial rate of photoentrainment while homeostatic adaptations in resilient-mice may lead to reduced light-induced SIK1 expression leading to faster photoentrainment.

The LHb exhibits independent oscillations in clock Per expression in the absence of SCN which highlights autonomous clock function in this nucleus that may drive certain circadian behaviours(3). Moreover, the neuropeptide prokineticin 2 (PK2) is part of the SCN output signal that conveys time of day information to the rest of the brain, including the LHb(33). Functional studies show that PK2 decreases LHb firing by increasing GABAergic tone(34). Thus, our observation of elevated day and night firing, together with delayed initial rate of photoentrainment, in the susceptible mice be indicative of decreased, and blunted rhythmic, PK2 signaling from the SCN. Furthermore, observations that the light pulse leads to increased firing in a subset of cells in the LHb(34) together with observation that hyperactivation of LHb cells leads to sleep disturbances possibly via DRN cells(35) highlights another putative mechanisms by which elevated firing in cells^DRN→SCN^ may leads to slow rate of photoentrainment in susceptible mice. Excitatory cortical projections to the DRN of susceptible-mice exhibit increased firing while feed-forward inhibition decreases firing in DRN cells(11). Thus, our current findings of decreased diurnal firing in DRN cells of susceptible-mice may be indicative of robust feedforward inhibition due to hyperactive cells^LHb→DRN^ in both the day and the night. In contrast, high day-time firing in the DRN of control and resilient-mice may be indicative of low LHb firing while low night-time firing in the DRN may be indicative of higher LHb firing. DRN’s direct and indirect projections to the SCN (cells^DRN→SCN^) regulates non-photic entrainment that modulate behaviours such as food gathering and social interactions(36, 37). In light of observations that decreased VTA dopamine input to the SCN slows the rate of photoentrainment(38), decreased diurnal firing in cells^DRN→SCN^ may be responsible for the delayed initial rate of photoentrainment in susceptible-mice. Moreover, our observations that the resilient-mice are able to photoentrainment much faster may be indicative of homeostatic adaptations resulting in large increases in day-time firing that we observed in DRN cells, a subset of which may be projecting to the SCN. These putative cells^DRN→SCN^ may modulate SCN cells such that the clock system is able to rapidly entrain to the new photo-stimulus.

We show that susceptible-mice exhibit: (i) pathophysiological changes in cells^LHb→DRN^ that lead to elevated diurnal firing, leading to blunted rhythmic activity and (ii) slower rate of photoentrainment. We speculate that, susceptible-mice may express dysfunctional cellular homeostatic mechanisms that lead to abnormal circadian clock gene rhythms that alters firing in circuits that regulate mood and photoentrainment. In contrast homeostatic adaptive changes in resilient-mice may buffer against pathophysiological changes. These studies continue to deepen our understanding of the link between stress and biological rhythms.

## Materials and Methods

### Animals

All experiments performed were approved by the NYUAD Animal Care and Use Committee, and all experimental protocols were conducted according to the National Institute of Health Guide for Care and Use of Laboratory Animals (IACUC Protocol: 150005A2). CD1 retired male breeders (Charles River), and C57BL/6J male mice (8-12 weeks; Jackson Laboratories) were used in all experiments of this study. All mice were maintained in the home cages, with *ad libitum* access (unless noted otherwise) to food and water in temperature (23 ± 2 °C)- and humidity (50 ± 10%)-controlled facilities with 12-hr light-dark (L/D) cycles (lights on: 7:00 AM; lights off: 7:00 PM, *Zeitgeber time ZT*; ZT0 - lights on; ZT12 - lights off). In phase-shift experiments mice were first entrained to the standard LD cycle after which they were exposed to a 6-hr phase advance where lights came on at 1:00 AM and lights off at 1:00 PM. All behavioral tests were conducted during the light cycle (ZT 5-10), and mice were habituated to the recording room and lighting conditions for at least 1-hr. Between trials, the behavioral apparatus was cleaned with MB-10 solution (Quip Laboratories, Inc. USA) to avoid persistence of olfactory cues.

### Viral vectors

AAV5-EF1a-DIO-mCherry, and AAV5-EF1a-DIO-hChR2 (H134R)-mCherry-WPRE-pA virus plasmids were purchased from the University of North Carolina vector core facility (UNC). Retrograde AAV5-CMV-PI-Cre-rBG virus was purchased from the University of Pennsylvania viral vector core facility (UPenn).

### Stereotaxic surgery, viral mediated gene transfer, and optic-fiber placement

Mice were anaesthetized with a ketamine (100 mg kg^−1^) and xylaxine (10 mg kg^−1^) mixture, placed in a stereotaxic apparatus (RWD Life Science Co., Ltd) and their skull was exposed by scalpel incision. For injection of replication defective viral vectors, 33-gauge needles were used. For injection of Cre-dependent AAV5-EF1a-DIO-mCherry virus, needles were placed bilaterally at a 0° angle into the LHb (in mm: anterior/posterior, –1.6; lateral/medial, 0.4; dorsal/ventral, –3.0) and 0.2 μl of virus was infused at a rate of 0.1 μl min^−1^. AAV5-EF1a-DIO-hChR2(H134R)-mCherry-WPRE-pA virus was injected bilaterally into the LHb at a 15° angle (in mm: anterior/posterior, –1.6; lateral/medial,1.1; dorsal/ventral, –3.1) and 0.2μl of virus was infused at a rate of 0.1 μl min^−1^. For retrograde travelling AAV5-CMV-PI-Cre-rBG viral injection, the needle was placed unilaterally at a 20° angle into the DRN (in mm: anterior/posterior, –4.7; lateral/medial, 1.1; dorsal/ventral, –3.2) and 0.5μl of virus was infused at a rate of 0.1 μl min^−1^. The needles were left in place for 10 min following injections to minimize diffusion and then completely withdrawn after viral delivery. For *in vivo* optical control of LHb neuronal firing in cells expressing hChR2-mCherry or mCherry, we used the chronically implantable optical-fiber system. Three weeks after surgery, chronically implantable homemade fibers (200 μm core optic fiber) were implanted above the LHb at a 15° angle (anterior–posterior, –1.6 mm; lateral–medial, 1.1 mm; dorsal–ventral, –3.0 mm). In order to ensure secure fixture of the implantable fiber, the skull was dried and then industrial strength dental cement (Zinc polycarboxylate cement; Factory of oral medicine materials at Stomatology School of Wuhan University) was added between the base of the implantable fiber and the skull. Mice were allowed to recover for at least 7-days before starting the behavioral paradigm.

### Blue light stimulation

Optical fibers (Thor Labs, BFL37-200) were connected using an FC/PC adaptor to a 473 nm blue laser diode (SLOC Lasers, BL473T8-150FC), and a stimulator (Agilent Technologies, no. 33220A) was used to generate blue light pulses. For *in vitro* slice electrophysiological validation of ChR2 activation, we tested 0.1–50 Hz (20 ms) stimulation protocols delivered to LHb neurons expressing ChR2 through an optic fiber (Thor Labs, BFL37-105) attached to a 473 nm laser. For all *in vivo* behavioral experiments, we injected mice with a cre-dependent AAV5-EF1a-DIO-mCherry or AAV5-EF1a-DIO-hChR2(H134R)-mCherry-WPRE-pA into LHb and replication defective retrograde travelling AAV5-CMV-PI-Cre-rBG into the DRN to specifically label the LHb→DRN circuit, and chronically implantable home-made fibers with 200 mm core optic fiber were implanted into the LHb as described above. All mice were handled for a minimum of 1-min per day for 4-days. Three days before the experiment, ‘dummy’ optical patch cables were connected to the fibers implanted in mice each day for 10 min in order to habituate them to the tethering procedure. During 7 days of subthreshold social defeat, mice received bilateral stimulation immediately after 10 min of physical interaction: mice were given >1-mW laser with a stimulation frequency of 20Hz and a 20ms width light pulse (500ms optical stimulation with an interstimulus interval of 1s) for 20-min/day during sensory contact.

### Chronic social defeat stress (CSDS) paradigm and social interaction (SI) test

Both chronic social defeat stress paradigm or subthreshold social-defeat stress paradigm were performed according to previously published protocols(14, 17, 18, 39, 40) with minor modifications in our lab. Briefly, CD1 aggressor mice were housed in social defeat cages 24 hrs before the onset of defeats on one side of a clear perforated plexiglass divider. The experimental C57 mice were individually exposed to an aggressive CD1 mouse for 10 min during which time they were physically attacked by the CD1 mouse. After 10 min of physical contact, the mice were separated by the clear perforated plexiglass divider. For the following 24 hrs, the aggressor and experimental mouse were maintained in sensory contact using the perforated plexiglass partition dividing the resident home cage in two. To avoid habituation, the experimental mice were exposed to a new CD1 aggressor mouse home cage each day for 15 consecutive days. The control mice were housed in pairs within a cage continuously separated by a clear perforated plexiglass divider. On the day following the last day of defeat, the mice were singly housed in new cages. Social interaction (SI) tests were performed on day 16. For the subthreshold social defeat stress (SSDS) paradigm, the procedure was identical to the standard CSDS paradigm, with the exception that the procedure lasted for 7 consecutive days. In order to stimulate LHb cells projecting to dorsal raphe nucleus (DRN), 200 mm core optic fibers were attached to the chronically implanted fibers, after which the experimental mice expressing hChR2-mCherry or mCherry in LHb-DRN circuit underwent bilateral blue light stimulation (20 min/day per hemisphere) during the sensory-stress period, immediately after 10 min of physical contact. The standard social defeat paradigm was carried out between 16:00 - 17:00, and subthreshold social defeat were performed between 13:00 - 17:00. Social-avoidance behaviour towards a novel CD1 non-aggressive mouse was measured in a two-stage SI test. In the first 2.5 min test (CD1 social target absent in the interaction zone), the experimental mouse was allowed to freely explore a square-shaped arena (44 × 44 cm) containing Plexiglas perforated cage with a wire mesh (10 × 6 cm) placed on one side of the arena. In the second 2.5 min test, the experimental mouse was reintroduced back into the arena with an unfamiliar CD1 non-aggressive mouse contained behind a Plexiglas perforated cage. TopScan video tracking system (CleverSys. Inc.) was used to automatically monitor and record the amount of time the experimental mouse spent in the ‘interaction zone’ (14 × 26 cm), ‘corner zone’ (10 × 10 cm) and ‘total travel’ within the arena for the duration of the 2.5 min test session (in absence and presence of the social target). Interaction zone time, corner zone time, total distance travelled and velocity were collected and analyzed. The segregation of susceptible and resilient mice was based on the social interaction ratio (SI ratio), which was calculated as [100 × (time spent in the interaction zone during social target present session)/ (time spent in the interaction zone during no social target session)] as described previously. All mice with scores < 100 were classified as ‘susceptible’ and those with scores ≥ 100 were classified as ‘resilient’.

### Sucrose preference test

To measure anhedonia, sucrose preference test was performed as we previously reported(28, 41, 42). After completion of the SI test, animals were single housed and habituated to two bottles of 1% sucrose for two days, followed by 24 hrs water and food deprivation. In the 3 hrs test period, the animals were exposed to one bottle of 1% sucrose and one bottle of water, with bottle positions switched half way through the experiment to ensure that the mice did not develop a side preference. The sucrose and water bottles were weighed before and after the test, recording the total consumption of each liquid. Sucrose preference was calculated as a percentage [100 × amount of sucrose consumption/total amount consumption (water + sucrose)].

### Circadian rhythms assay

Circadian activity rhythms of mice were determined in individual cages (33.2 × 15 × 13 cm) equipped with stainless steel wheels (11 cm inside diameter, 5.4 cm wide; Model ACTI-PT2-MCR2, Actimetrics, IL, USA). Wheel-running cages were placed in circadian cabinets (Phenome Technologies, Inc., IL, USA) with 6 cages per row. The light and temperature of the chambers was controlled by the ClockLab Chamber Control software (ACT-500, Actimetrics, IL, USA). Circadian activity rhythms were measured with infrared (clickless) sensor clips onto the lip and rail of the cage that detected wheel rotations, and the number of wheel revolutions every minute. Wheel rotation from the sensor was transmitted via a single channel connected to the ClockLab digital interface (ACT-556, Actimetrics, IL, USA) and recorded by a personal computer. Data were collected and analyzed using ClockLab Data Collection software (ACT-500, Actimetrics, IL, USA). All data was analyzed by a trained scorer blind to treatment. To record the circadian activity rhythms, mice were individually housed in activity wheel-equipped cages (Actimetrics, IL, USA) in light-tight boxes under a 12-h light–dark cycle. (light intensity 180–200 lux; temperature inside light-tight boxes: 25.5 ± 1.5°C) for at least 7 days, then transferred to constant and complete darkness (DD) cycle and provided with food and water *ad libitum*. Wheel running rhythms were monitored and analyzed with ClockLab Data Collection system (ACT-500, Actimetrics, IL, USA). The free-running period was calculated according to the onset of activity across seven days in constant darkness. Onset of activity was identified as the first bin above a threshold of 5 counts preceded by at least 6 hrs of inactivity and followed by at least 6 hrs of activity and offsets were determined by at least 6 hrs of activity and followed by 6 hrs of inactivity through ClockLab software. Onset and offset points were edited by eye when necessary. A regression line was fit through activity onsets for the 7 days of LD cycle and extrapolated for 7 days following DD cycle. The magnitude of phase shifts was calculated as the time difference between the two lines of LD and DD cycle. To assess re-entrainment to a 6-h advance to the LD cycle, mice were kept in a 12-h light–dark cycle (light intensity 180–200 lux; temperature inside light-tight boxes: 25.5 ± 1.5°C) for at least 7 days. Following this entrainment, the onset of darkness of the cycle was abruptly advanced 6 hrs and daily advances (hours) in running wheel activity onsets were recorded every day for an additional 7 days. The days required to entrain following the shift in the light cycle was calculated. The duration of re-entrainment was defined as the number of days required to shift activity onset by 6 ± 0.25 hrs followed by two consecutive days of activity onset within this range.

### In vitro patch-clamp electrophysiology

Cell-attached or whole-cell recordings were obtained from LHb neurons projecting to DRN in acute brain slices from C57BL/6J mice that had been stereotaxically injected with AAV5-EF1a-DIO-mCherry or AAV5-EF1a-DIO-hChR2(H134R)-mCherry-WPRE-pA into the LHb, and retrograde viral vector AAV5-CMV-PI-Cre-rBG in DRN. To minimize stress and to obtain healthy LHb slices, mice were anaesthetized with isofluorane and perfused immediately for 40–60 s with ice-cold artificial cerebrospinal fluid (aCSF), which contained (in mM): 128 NaCl, 3 KCl, 1.25 NaH2PO4, 10 D-glucose, 24 NaHCO3, 2 CaCl2 and 2 MgCl2 (oxygenated with 95% O2 and 5% CO2, pH 7.4, 295–305 mOsm). Acute brain slices containing LHb neurons were cut using a microslicer (DTK-1000, Ted Pella) in ice-cold sucrose aCSF, which was derived by fully replacing NaCl with 254 mM sucrose and saturated by 95% O2 and 5% CO2. Slices were maintained in holding chambers with aCSF for 1 hr recovery at 37 °C and then at room temperature for recording. Cell-attached and whole-cell recordings were carried out using aCSF at 34°C (flow rate = 2.5 ml min^−1^). Patch pipettes (5-8 MΩ for cell-attached recordings, and 3-5 MΩ for whole-cell recordings) were filled with internal solution containing the following (in mM): 115 potassium gluconate, 20 KCl, 1.5 MgCl2, 10 phosphocreatine, 10 HEPES, 2 magnesium ATP and 0.5 GTP (pH 7.2, 285 mOsm). For measurements of the spontaneous activity of mCherry-labeled LHb-DRN neurons, cell-attached recordings were performed in acutely-prepared LHb-containing brain slices. Signals were band-pass filtered at 300 Hz – 1k Hz and then Bessel filtered at 10 kHz (gain 1) using a Multiclamp 700B amplifier. To measure the intrinsic membrane properties of LHb neurons, whole-cell recordings were carried out in current-clamp mode and spikes were induced by incremental increases of current injection (each step increase was 25 pA; range -10–200 pA). Whole-cell voltage clamp was used to record Ih-current with a series of 2 s pulses with a 10mV command voltage step from −120 mV to −60 mV from a holding potential of −60 mV. To isolate voltage-gated K^+^ channel mediated currents, 3 s pulses with a 10 mV from 0 mV to +80 mV from a holding potential of −60 mV.

For whole-cell recordings during optogenetic stimulation, resting membrane potential and action potentials were recorded in current-clamp mode and inward current measurements were made in voltage-clamp mode using the Multiclamp 700B amplifier and data acquisition was collected using a Digidata 1550B digitizer and Clampfit 10.5 (Molecular Devices). Series resistance was monitored during the experiments and membrane currents and voltages were filtered at 10 kHz (Bessel filter), and cells were discarded from further analysis if under their basal conditions this value changed by more than 15%. For *in vitro* electrophysiological validation of ChR2 activation, whole cell optogenetic recordings were obtained from LHb neurons in acute brain slices from C57BL/6J mice that had been stereotaxically injected with AAV5-EF1a-DIO-hChR2(H134R)-mCherry-WPRE-pA into the LHb, and retrograde viral vector AAV5-CMV-PI-Cre-rBG in DRN. Sustained and trains (0.1–50 Hz) of blue light were generated by a stimulator (described above) and delivered to LHb neurons expressing ChR2 through a 105 mm optic fiber attached to a 473 nm laser. Day-time recordings were performed at ZT2-7 while night-time recordings were performed at ZT14-19.

### In vivo extracellular single unit electrophysiology

Mice were anesthetized with 10% chloral hydrate (400mg/Kg), and head fixed horizontally onto a stereotaxic frame. Using bregma, LHb was located within the range (in mm): anterior/posterior: −0.94 to −2.18, medial/lateral: 0.25 to 0.80, dorsal/ventral: −2.50 to −2.80. Glass micropipettes (15-20 MΩ) filled with 2M NaCl was used for recording. Recorded electrical signals were amplified and band-pass filtered (0.3k-1k Hz) using a DP-311 Differential Amplifier (Warner Instruments). Data acquisition of *in vivo* spontaneous firing activity was collected using a Digidata 1550B digitizer and Clampfit 10.5 (Molecular Devices).

### Immunohistochemistry

Mice were transcardially perfused with 20 ml saline, followed by 20 ml 4% (weight divided by volume; wt/vol) phosphate buffer saline (PBS)-buffered paraformaldehyde under deep anesthesia with 10% chloral hydrate (400 mg/Kg). Brains were dissected and post-fixed overnight at 4 °C, then treated with 30% sucrose at 4 °C for 2 days. Coronal sections with thickness of 30 μm were cut with Leica VT1200 Semiautomatic Vibrating Blade Microtome. Sections were permeabilized with PBS-T (0.1% Triton 100X, Sigma) and blocked in PBS-T including 5% (wt/vol) normal donkey serum (Sigma) for one hour. For labelling neurons, the sections were incubated in the primary antibody rabbit anti-NeuN (1:500; 26975-I-AP, ProteinTech) diluted in PBS-T with 5% normal donkey serum at 4 °C with gentle shaking for three days, then washed three times in PBS-T. For visualization, anti-NeuN was detected by the secondary antibody goat anti-Rabbit Alexa 488 (1:500; 711-545-152, Jackson ImmunoResearch Laboratories), and all sections were counter-stained with DAPI (300 nM; final concentration Sigma) for two days’ staining at 4°C with gentle shaking. Sections were then washed and mounted in PBS on Polysine microscope slides (VWR International, LLC.), air-dried overnight and coverslipped with Aquatex aqueous mounting agent (VWR International, LLC.). All images were captured on Olympus FV1000 confocal microscope (Olympus Corporation, Japan) at 10X magnification at the New York University Abu Dhabi Microscope Core facility and analyzed with Olympus FV1000 software.

### Statistical analysis

The estimated sample sizes were based on our past experience performing similar experiments and power analysis. Animals were randomly assigned to treatment groups. Analyses were performed in a manner blinded to treatment assignments in all behavioral experiments. Data are presented as mean ± SEM. All analysis was performed using GraphPad Prism software v7 for Windows (La Jolla, California). By pre-established criteria, values were excluded from analyses if the viral injection sites were out of the LHb. When comparing two groups of normally distributed data, unpaired student’s two tailed t-test was used. Multiple-groups comparisons were achieved by means of analysis of variance with/without repeated factors (one-way or two-way ANOVA), followed by a post hoc Dunnett’s/Tukey’s multiple comparisons test. Two criteria were used to confirm burst firing: (1) Onset was defined as spikes beginning with a maximal inter-spike interval of 20 ms and ended with a maximal inter-spike interval of 100 ms, and minimum intra-burst interval is 100 ms and (2) minimum number of spikes in a burst is 2. All statistical tests were two-tailed, and statistical significance was set at P < 0.05.

## ACKNOWLEDGEMENTS AND DISCLOSURES

This study was supported in part by grants from (NYUAD Annual Research Budget, Brain and Behavior Research Foundation (NARSAD), NYUAD Research Enhancement Fund, NYU Research Enhancement Fund, Al Jalila Research Foundation: DC), the National Natural Science Foundation of China (NSFC81300957: HL), the Natural Science Foundation of Jiangsu Province (BK20181145: HL), the Research Start-up Funding for Talent Introduction (2019203002: HL), and the Clinical Technical Research and Study Plan Project (2018211006: HL). The authors declare no competing financial interests.

---

**Suppl. Figure 1.**
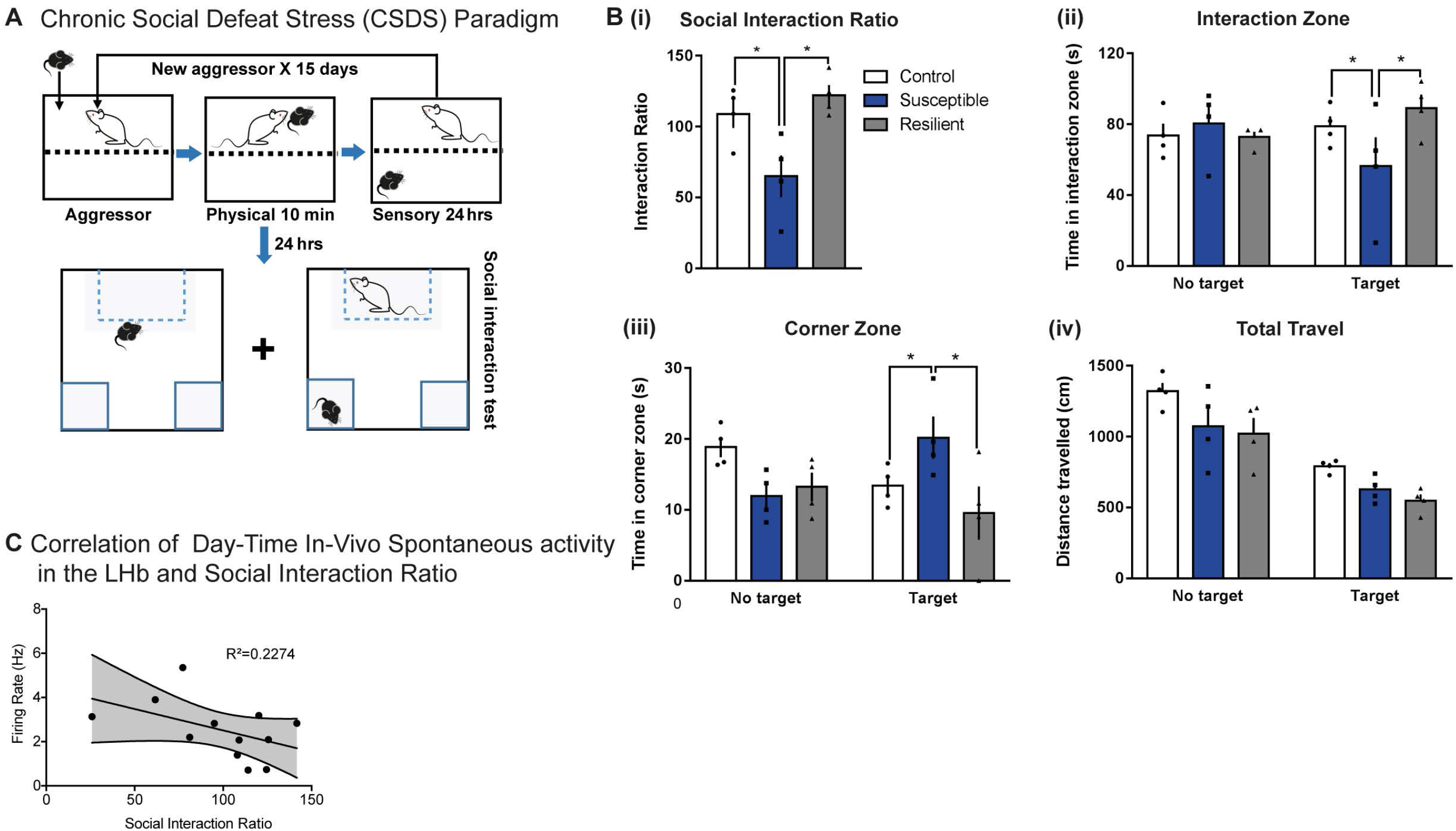
Social Stress Induces Social Avoidance in Susceptible-mice. **(A)** Detailed schematic of the 15-day CSDS paradigm using C57BL/6J male mice and CD1 retired male breeders (aggressor) and SI test. **(B)** SI data showed that in the presence of a CD1 social target (non-aggressor), susceptible mice displaying **(i)** decreased social interaction ratio (*F*_2,9_ = 7.295, *P* < 0.05; n = 4 mice/group), **(ii)** decreased time in the interaction zone (*F*_2,9_ = 5.185, *P* < 0.05), **(iii)** increased time in the corner zone (*F*_2,9_ = 5.188, *P* < 0.05) and **(iv)** no difference in total travel between control, susceptible and resilient mice. **(C)** Though not significant, correlation analysis of in-vivo spontaneous activity in the LHb and social interaction ratio showed a very slight negative correlation. Error bars: mean ± SEM.

**Suppl. Figure 2.**
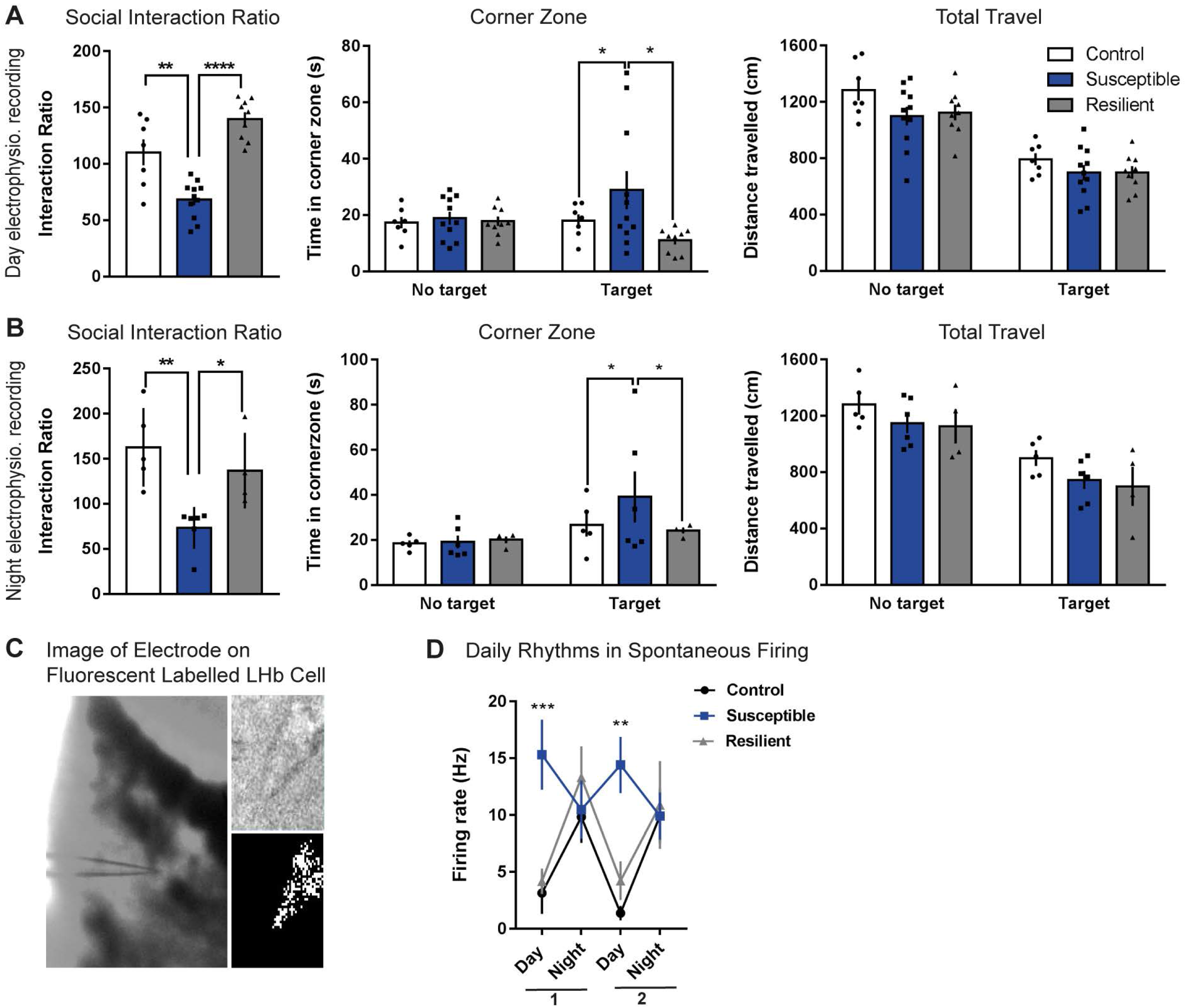
Social Interaction Data in Mice Used to Measure Day or Night Time Spontaneous Firing in Cells^LHb→DRN^. **(A)** Susceptible-mice used for day- or, **(B)** night-time *in vitro* recording of labelled cells^LHb-DRN^ display: decreased social interaction ratio (day - *F*_2,24_ = 28.02, *P* < 0.0001; n = 7-11 mice/group; night - *F*_2,12_ = 8.959, *P* < 0.01; n = 4-6 mice/group) (left). In the presence of CD1 (social target) susceptible-mice spent increased time in the corner zone (day - *F*_2,24_ = 3.335, *P* < 0.05; night - *F*_2,12_ = 6.444, *P* < 0.05) (middle). There was no difference in total travel between control, susceptible and resilient-mice (right). **(C)** Representative image of electrode on a fluorescent labelled cell^LHb-DRN^. **(D)** The *in-vitro* electrophysiology experiments were performed over 16-days after the SI test. To better visualize the rhythmic changes in spontaneous firing in cells^LHb→DRN^, data was re-graphed where day and night firing was binned into 1^st^ half (1-8 Day/Night after SI) and the 2^nd^ half (9-16 Day/Night after SI) recording sessions. In susceptible-mice, the day time firing pattern is phase inversed and was significantly higher on day 1 than control and resilient mice (Day 1 – control vs susceptible: *F*_2,160_ = 3.617, *P* = 0.0306; resilient vs susceptible: *F*_2,160_ = 3.689, *P* = 0.0268; n = 53-64 cells from 7 to 11 mice/group), and day 2 from control mice (Day 2 – *F*_2,160_ = 3.653, *P* = 0.0286), *** - day vs day in control vs susceptible and resilient vs susceptible mice, ** - control vs susceptible. Error bars: mean ± SEM

**Supplementary Figure 3.**
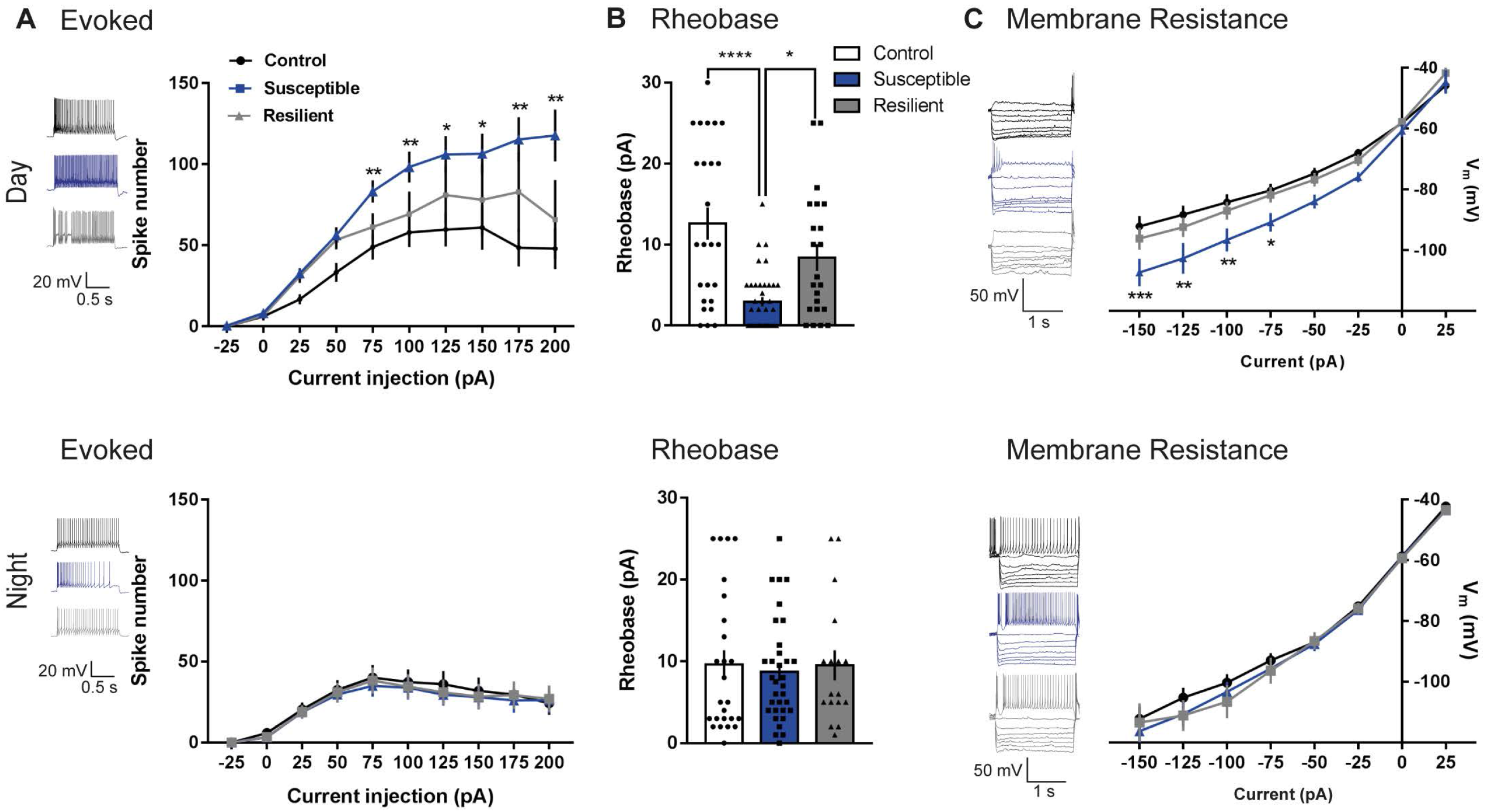
Diurnal Difference in Intrinsic Membrane Properties in Mice Exposed to CSDS. **(A)** Sample traces of evoked firings of cells^LHb-DRN^ in day (top left) or night (bottom left) in control (top), susceptible (middle) and resilient-mice (bottom). cells^LHb-DRN^ from susceptible-mice display increased day-time excitability in response to incremental steps in current injections (75, 100, 125, 150, 175 and 200 pA) compared with control and resilient-mice (*F*_2,820_ = 21.61, *P* < 0.0001; n = 14 to 42 cells from 7 to 11 mice per group) (top right). There was no difference between the stress phenotypes at night (bottom right). **(B)** Cells^LHb-DRN^ from susceptible-mice display decreased day-time rheobase (*F*_2,82_ = 14.37, *P* < 0.0001; n = 22 to 38 cells from 7 to 11 mice per group) (top). There was no difference in night-time rheobase between the stress phenotypes (bottom). **(C)** Representative voltage traces in response to current injections in cells^LHb-DRN^ in day (top left) or night (bottom left) in control (top), susceptible (middle) and resilient-mice (bottom). Day-time *I–V* relationship showing cells^LHb-DRN^ from susceptible-mice display increased membrane resistance (*F*_2,264_ = 23.13, *P* < 0.0001; n = 9 to 15 cells from 7 to 11 mice per group) (top right). There was no difference in Night-time *I–V* relationship between the stress phenotypes (bottom right). Error bars: mean ± SEM.

**Suppl. Figure 4.**
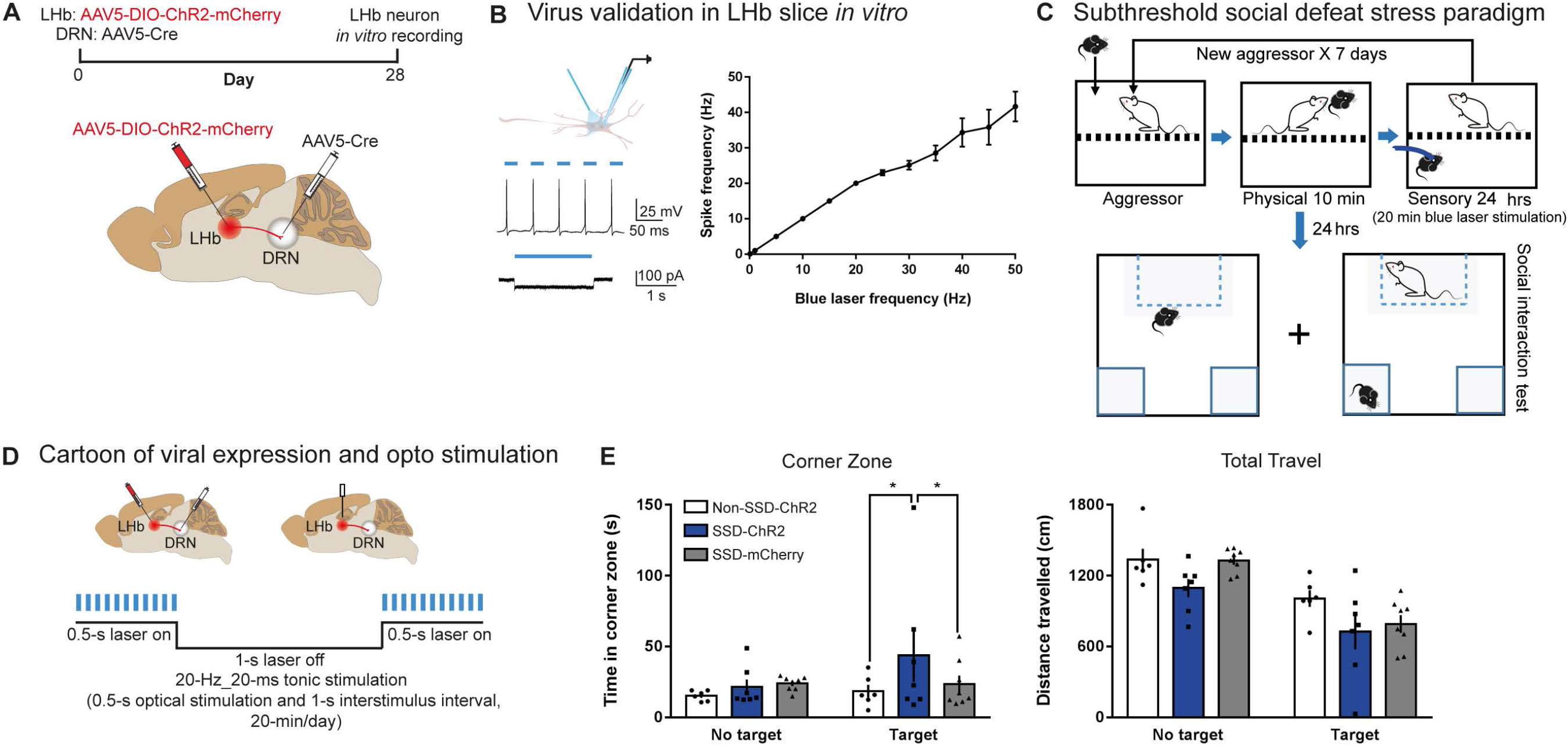
Functional validation of AAV-DIO-ChR2-mCherry and optical stimulation after each exposure to social stress for 7 days SSDS paradigm. **(A)** Experimental timeline of pathway-specific *in-vitro* recording of labelled cells^LHb-DRN^ (top), and schematic showing surgeries site for virus injections to specifically label cells^LHb-DRN^ (bottom). (**B)** Schematic showing the optical fiber used for *in-vitro* delivery of blue light (470 nm) and glass electrode used for simultaneous extracellular recording of labelled cells^LHb-DRN^ responses (top left). Whole-cell current-clamp recordings from AAV-DIO-ChR2-mCherry infected cells^LHb-DRN^ in LHb slices showing 20 Hz_20 ms of blue light stimulation induces tonic firings (middle left). Whole-cell voltage clamp recordings showing long duration (2s) of blue light stimulation induces temporally precise inward photocurrent (bottom left), Frequency-response curve of membrane excitability to blue light stimulation showing light frequencies ranging from 0.1-20 Hz reliably induce the approximately equivalent firing rate in the ChR2-mCherry expressing cells^LHb-DRN^ (right). **(C)** Detailed schematic of the 7-day SSDS procedure during which the ChR2 expressing cells^LHb-DRN^ were optically stimulated for 20-mins after each exposure to social stress for 7 days, after which the mice underwent the SI test. **(D)** Cartoon showing *in-vivo* optical stimulation protocol during the SSDS paradigm. **(E)** In the presence of a CD1 social target (non-aggressor), SSDS-ChR2 mice display increased time in the corner zone (*F*_2,18_ = 3.55, *P* < 0.05) (right). There was no difference in total travel (right). Error bars: mean ± SEM.

**Suppl. Figure 5.**
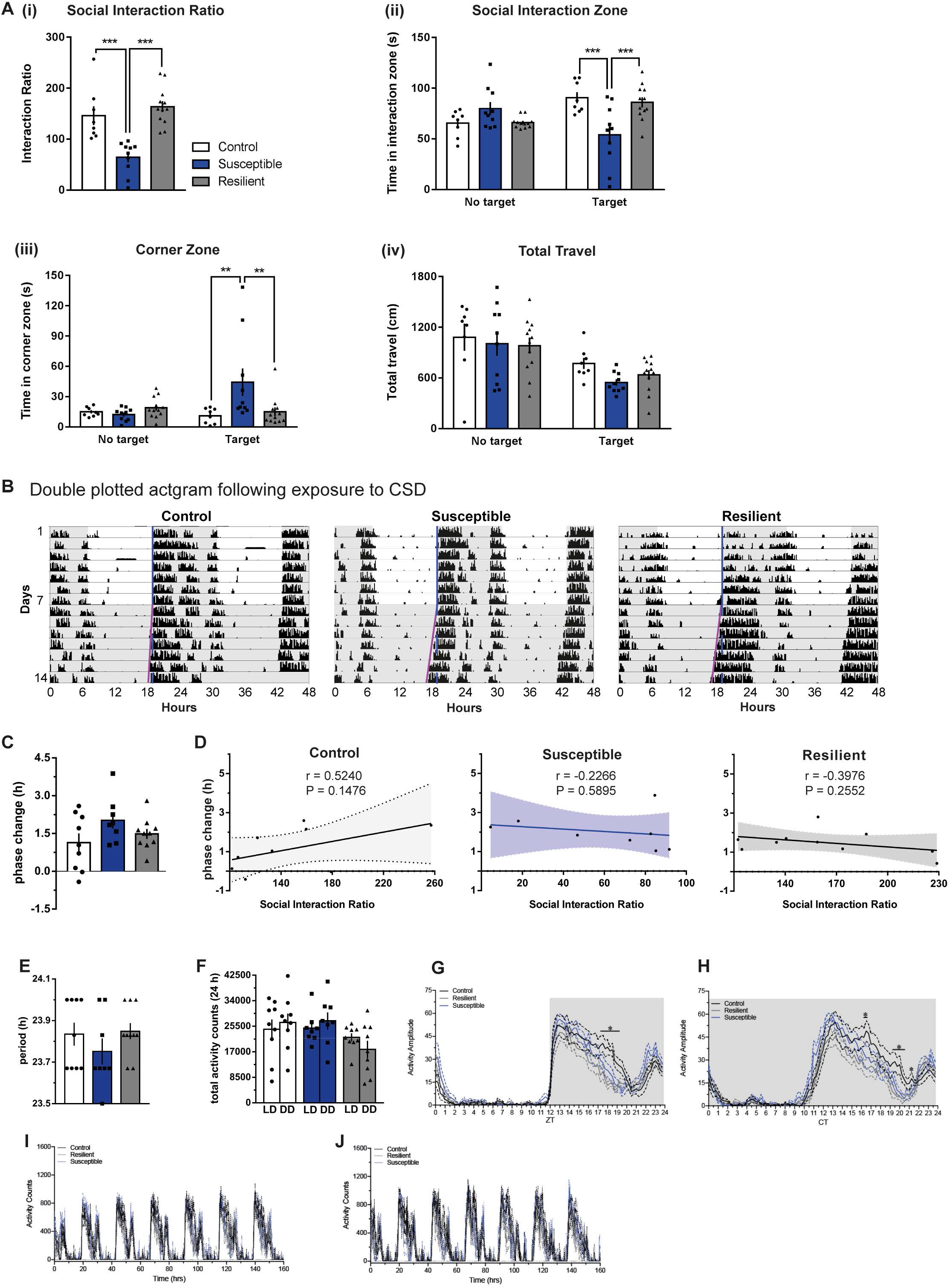
No Difference in Endogenous Rhythms in Mice Exposed to Chronic Social Stress. **(A)** In the presence of a CD1 social target (non-aggressor), susceptible-mice displaying **(i)** decreased social interaction ratio (*F*_2,27_ = 18.08, *P* < 0.0001; n = 8-12 mice/group), **(ii)** decreased time in the interaction zone (*F*_2,27_ = 19.68, *P* < 0.0001), **(iii)** increased time in the corner zone (*F*_2,27_ = 5.188, *P* < 0.05), **(iv)** no difference in total travel. **(B)** Representative double-plotted actograms of control-, susceptible- and resilient-mice. Gray background indicating either dark phase of the LD cycle or constant darkness (DD); light blue and pink line representing extended regression line derived by onset of activity under LD and DD cycle, respectively. **(C-F)** No difference in - (c) phase change in activity onset in DD, - (d) in correlation between interaction ratio and degree of phase change, - (e) in free running period length in DD, and - (f) in total activity counts in 24-hrs between LD and DD between control, susceptible and resilient-mice. Though not significant, susceptible mice exhibit slightly larger phase change (c) and shorter free running period (e). Error bars: mean ± SEM.

**Suppl. Figure 6.**
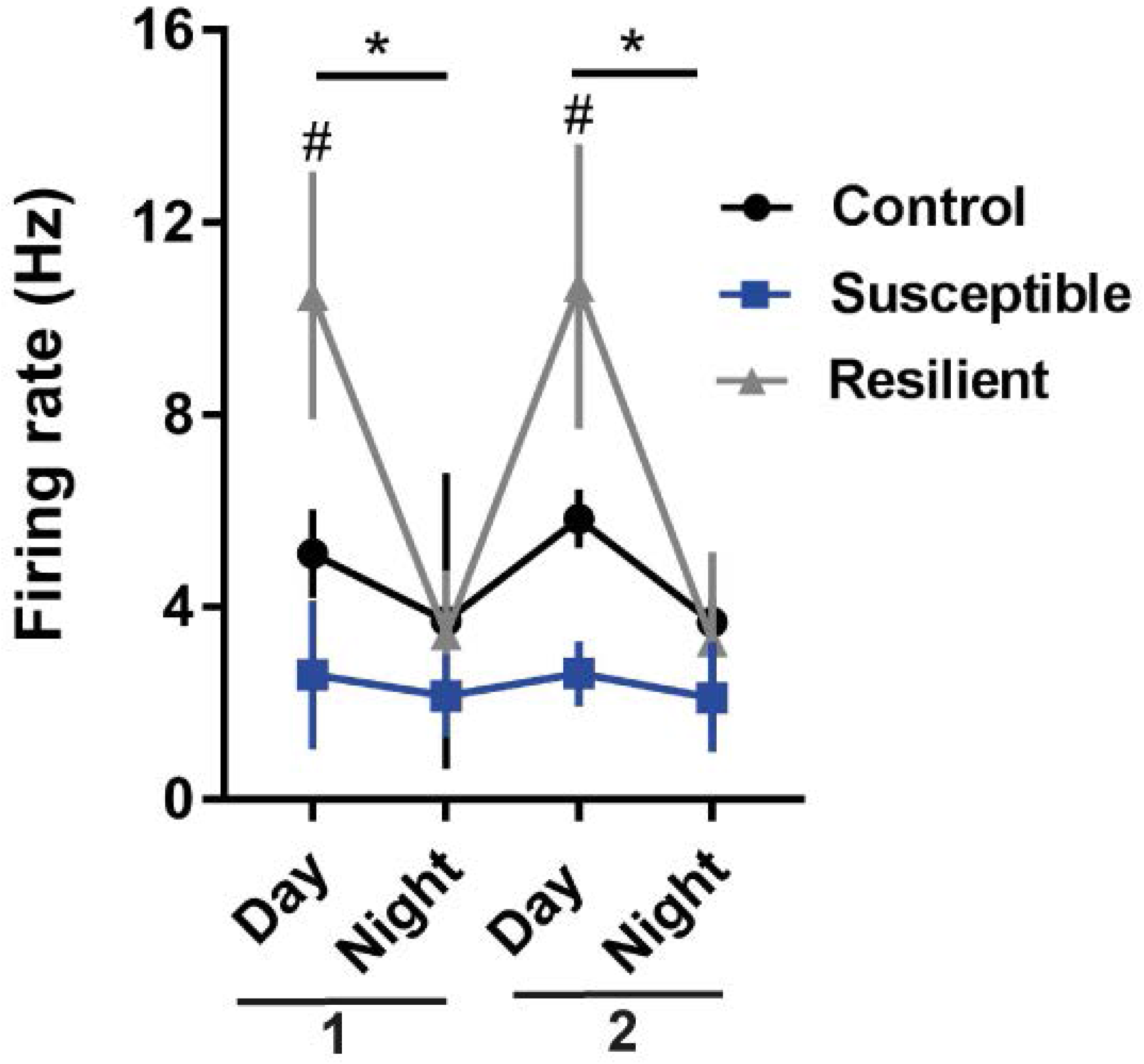
Blunted Rhythms in Spontaneous Firing in DRN Cells of Stress-Susceptible Mice. **(A)** The *in-vitro* electrophysiology experiments were performed over 16-days after the SI test. To better visualize the rhythmic changes in spontaneous firing in cells^LHb→DRN^, data was re-graphed where day and night firing was binned into 1^st^ half (1-8 Day/Night after SI) and the 2^nd^ half (9-16 Day/Night after SI) recording sessions. DRN cells display daily rhythms in spontaneous firings in resilient-mice with significantly higher daytime activity than night (Day 1 - *F*_2,79_ = 4.125, *P* = 0.0234; Day 2 - *F*_2,79_ = 4.111, *P* = 0.0240; n = 53-64 cells from 4 to 10 mice/group). Also, in daytime, spontaneous firing of DRN cells was significantly higher in resilient-mice than susceptible (Day 1 - *F*_3,79_ = 4.085, *P* = 0.0137; Day 2 - *F*_3,79_ = 4.072, *P* = 0.0141; n = 53-64 cells from 4 to 10 mice/group). * - day vs night in resilient mice, #- day vs day in resilient vs susceptible mice, Error bars: mean ± SEM.

## Reference

1. Cho JR, Treweek JB, Robinson JE, Xiao C, Bremner LR, Greenbaum A, et al. Dorsal Raphe Dopamine Neurons Modulate Arousal and Promote Wakefulness by Salient Stimuli. Neuron. 2017;94(6):1205–19 e8.

2. Zhao H, Zhang BL, Yang SJ, Rusak B. The role of lateral habenula-dorsal raphe nucleus circuits in higher brain functions and psychiatric illness. Behav Brain Res. 2015; 277:89–98.

3. Salaberry NL, Hamm H, Felder-Schmittbuhl MP, Mendoza J. A suprachiasmatic-independent circadian clock(s) in the habenula is affected by Per gene mutations and housing light conditions in mice. Brain structure & function. 2019;224(1):19–31.

4. Bano-Otalora B, Piggins HD. Contributions of the lateral habenula to circadian timekeeping. Pharmacol Biochem Behav. 2017;162:46–54.

5. Souetre E, Salvati E, Belugou JL, Pringuey D, Candito M, Krebs B, et al. Circadian rhythms in depression and recovery: evidence for blunted amplitude as the main chronobiological abnormality. Psychiatry Res. 1989;28(3):263–78.

6. Bechtel W. Circadian Rhythms and Mood Disorders: Are the Phenomena and Mechanisms Causally Related? Frontiers in psychiatry. 2015;6:118.

7. Hikosaka O. The habenula: from stress evasion to value-based decision-making. Nat Rev Neurosci. 2010;11(7):503–13.

8. Zhao H, Rusak B. Circadian firing-rate rhythms and light responses of rat habenular nucleus neurons in vivo and in vitro. Neuroscience. 2005;132(2):519–28.

9. Guilding C, Hughes AT, Piggins HD. Circadian oscillators in the epithalamus. Neuroscience. 2010;169(4):1630–9.

10. Monti JM. The role of dorsal raphe nucleus serotonergic and non-serotonergic neurons, and of their receptors, in regulating waking and rapid eye movement (REM) sleep. Sleep Med Rev. 2010;14(5):319–27.

11. Challis C, Boulden J, Veerakumar A, Espallergues J, Vassoler FM, Pierce RC, et al. Raphe GABAergic neurons mediate the acquisition of avoidance after social defeat. J Neurosci. 2013;33(35):13978–88,88a.

12. Kim U, Chang SY. Dendritic morphology, local circuitry, and intrinsic electrophysiology of neurons in the rat medial and lateral habenular nuclei of the epithalamus. J Comp Neurol. 2005;483(2):236–50.

13. Atkinson SE, Maywood ES, Chesham JE, Wozny C, Colwell CS, Hastings MH, et al. Cyclic AMP signaling control of action potential firing rate and molecular circadian pacemaking in the suprachiasmatic nucleus. J Biol Rhythms. 2011;26(3):210–20.

14. Friedman AK, Walsh JJ, Juarez B, Ku SM, Chaudhury D, Wang J, et al. Enhancing depression mechanisms in midbrain dopamine neurons achieves homeostatic resilience. Science. 2014;344(6181):313–9.

15. Itri JN, Vosko AM, Schroeder A, Dragich JM, Michel S, Colwell CS. Circadian regulation of a-type potassium currents in the suprachiasmatic nucleus. J Neurophysiol. 2010;103(2):632–40.

16. Kudo T, Loh DH, Kuljis D, Constance C, Colwell CS. Fast delayed rectifier potassium current: critical for input and output of the circadian system. J Neurosci. 2011;31(8):2746–55.

17. Chaudhury D, Walsh JJ, Friedman AK, Juarez B, Ku SM, Koo JW, et al. Rapid regulation of depression-related behaviours by control of midbrain dopamine neurons. Nature. 2013;493(7433):532–6.

18. Krishnan V, Han MH, Graham DL, Berton O, Renthal W, Russo SJ, et al. Molecular adaptations underlying susceptibility and resistance to social defeat in brain reward regions. Cell. 2007;131(2):391–404.

19. Zhang C, Truong KK, Zhou QY. Efferent projections of prokineticin 2 expressing neurons in the mouse suprachiasmatic nucleus. PLoS One. 2009;4(9):e7151.

20. Koch CE, Leinweber B, Drengberg BC, Blaum C, Oster H. Interaction between circadian rhythms and stress. Neurobiology of stress. 2017;6:57–67.

21. Nicolaides NC, Charmandari E, Kino T, Chrousos GP. Stress-Related and Circadian Secretion and Target Tissue Actions of Glucocorticoids: Impact on Health. Frontiers in endocrinology. 2017;8:70.

22. Christiansen SL, Bouzinova EV, Fahrenkrug J, Wiborg O. Altered Expression Pattern of Clock Genes in a Rat Model of Depression. Int J Neuropsychopharmacol. 2016.

23. Jacinto LR, Mata R, Novais A, Marques F, Sousa N. The habenula as a critical node in chronic stress-related anxiety. Exp Neurol. 2017;289:46–54.

24. Christiansen S, Bouzinova EV, Palme R, Wiborg O. Circadian activity of the hypothalamic-pituitary-adrenal axis is differentially affected in the rat chronic mild stress model of depression. Stress. 2012;15(6):647–57.

25. Jiang WG, Li SX, Zhou SJ, Sun Y, Shi J, Lu L. Chronic unpredictable stress induces a reversible change of PER2 rhythm in the suprachiasmatic nucleus. Brain Res. 2011;1399:25–32.

26. Kinoshita C, Miyazaki K, Ishida N. Chronic stress affects PERIOD2 expression through glycogen synthase kinase-3beta phosphorylation in the central clock. Neuroreport. 2012;23(2):98–102.

27. Colwell CS. Linking neural activity and molecular oscillations in the SCN. Nat Rev Neurosci. 2011;12(10):553–69.

28. Yang Y, Cui Y, Sang K, Dong Y, Ni Z, Ma S, et al. Ketamine blocks bursting in the lateral habenula to rapidly relieve depression. Nature. 2018;554(7692):317–22.

29. Wilkinson MB, Dias C, Magida J, Mazei-Robison M, Lobo M, Kennedy P, et al. A novel role of the WNT-dishevelled-GSK3beta signaling cascade in the mouse nucleus accumbens in a social defeat model of depression. J Neurosci. 2011;31(25):9084–92.

30. Paul JR, McKeown AS, Davis JA, Totsch SK, Mintz EM, Kraft TW, et al. Glycogen synthase kinase 3 regulates photic signaling in the suprachiasmatic nucleus. Eur J Neurosci. 2017;45(8):1102–10.

31. Jagannath A, Butler R, Godinho SIH, Couch Y, Brown LA, Vasudevan SR, et al. The CRTC1-SIK1 pathway regulates entrainment of the circadian clock. Cell. 2013;154(5):1100–11.

32. Liu Y, Poon V, Sanchez-Watts G, Watts AG, Takemori H, Aguilera G. Salt-inducible kinase is involved in the regulation of corticotropin-releasing hormone transcription in hypothalamic neurons in rats. Endocrinology. 2012;153(1):223–33.

33. Cheng MY, Bullock CM, Li C, Lee AG, Bermak JC, Belluzzi J, et al. Prokineticin 2 transmits the behavioural circadian rhythm of the suprachiasmatic nucleus. Nature. 2002;417(6887):405–10.

34. Sakhi K, Wegner S, Belle MD, Howarth M, Delagrange P, Brown TM, et al. Intrinsic and extrinsic cues regulate the daily profile of mouse lateral habenula neuronal activity. J Physiol. 2014;592(Pt 22):5025–45.

35. Aizawa H, Cui W, Tanaka K, Okamoto H. Hyperactivation of the habenula as a link between depression and sleep disturbance. Front Hum Neurosci. 2013;7:826.

36. Edgar DM, Reid MS, Dement WC. Serotonergic afferents mediate activity-dependent entrainment of the mouse circadian clock. Am J Physiol. 1997;273(1 Pt 2):R265–9.

37. Marchant EG, Watson NV, Mistlberger RE. Both neuropeptide Y and serotonin are necessary for entrainment of circadian rhythms in mice by daily treadmill running schedules. J Neurosci. 1997;17(20):7974–87.

38. Grippo RM, Purohit AM, Zhang Q, Zweifel LS, Guler AD. Direct Midbrain Dopamine Input to the Suprachiasmatic Nucleus Accelerates Circadian Entrainment. Curr Biol. 2017;27(16):2465–75 e3.

39. Berton O, McClung CA, Dileone RJ, Krishnan V, Renthal W, Russo SJ, et al. Essential role of BDNF in the mesolimbic dopamine pathway in social defeat stress. Science. 2006;311(5762):864–8.

40. Cao JL, Covington HE, 3rd, Friedman AK, Wilkinson MB, Walsh JJ, Cooper DC, et al. Mesolimbic dopamine neurons in the brain reward circuit mediate susceptibility to social defeat and antidepressant action. J Neurosci. 2010;30(49):16453–8.

41. Powell TR, Fernandes C, Schalkwyk LC. Depression-Related Behavioral Tests. Curr Protoc Mouse Biol. 2012;2(2):119–27.

42. Cui Y, Yang Y, Ni Z, Dong Y, Cai G, Foncelle A, et al. Astroglial Kir4.1 in the lateral habenula drives neuronal bursts in depression. Nature. 2018;554(7692):323–7.

